# Contribution of ammonia oxidizers to inorganic carbon fixation in the dark ocean

**DOI:** 10.1101/2024.11.16.623942

**Authors:** Barbara Bayer, Katharina Kitzinger, Nicola L. Paul, Justine B. Albers, Mak A. Saito, Michael Wagner, Craig A. Carlson, Alyson E. Santoro

## Abstract

Ammonia-oxidizing archaea are the most abundant chemolithoautotrophs in the ocean, comprising up to 40% of microbial cells in deep waters, and are assumed to dominate dissolved inorganic carbon (DIC) fixation below the sunlit surface layer. Yet, the supply of reduced nitrogen from particulate organic matter flux from the surface is insufficient to support the amount of nitrification required to sustain measured DIC fixation rates in the dark ocean. The aim of this study was to quantify the contribution of ammonia oxidizers to DIC fixation in the dark ocean. We used phenylacetylene - a specific inhibitor of the ammonia monooxygenase enzyme - to selectively inhibit ammonia oxidizers during two oceanographic expeditions in the eastern tropical and subtropical Pacific Ocean spanning 35º N to 10º S. We show that ammonia oxidizers contribute only a small fraction to dark DIC fixation, accounting for 2 to 22% of the depth-integrated rates in the eastern tropical Pacific. The highest contributions were observed at the depth of the nitrification maximum, where ammonia oxidation could account for up to 50% of dark DIC fixation. Our results help to reconcile the observed discrepancies between nitrogen supply and DIC fixation at depth, and provide a new perspective on global ocean chemolithoautotrophy, revealing that the majority of DIC fixation within the lower euphotic zone and below 200 m depth is not fueled by ammonia oxidation.

**Significance:** Microbes in the ocean play important roles in the global carbon cycle and the ocean’s capacity to sequester carbon. Despite this importance, deciphering the contributions of different microbial metabolic processes to the oceanic carbon budget remains challenging. Particularly in the dark ocean, large discrepancies between organic matter fluxes and measured microbial metabolic rates are observed. We show that abundant chemoautotrophs – ammonia-oxidizing microbes – contribute only a small fraction to dark carbon fixation in the Pacific Ocean, challenging the current view that carbon fixation in the dark ocean is primarily sustained by nitrification. This work advances our understanding of microbial carbon processing, and offers new insights into the long-standing question of the main energy sources fueling carbon fixation in the dark ocean.

## Introduction

Phytoplankton-driven ocean primary production is one of the most important biological processes converting dissolved inorganic carbon (DIC) into organic matter, forming the base of the marine food web (1). While most of the DIC fixation in the surface ocean is fueled by light energy, non-photosynthetic DIC fixation (= “dark DIC fixation”) has been proposed to contribute substantially to the biological uptake of DIC in the ocean (2–4), possibly increasing global ocean primary production estimates by up to 22% (5, 6). In surface waters, dark DIC fixation has been primarily associated with the activities of heterotrophic bacteria (7, 8), which can incorporate DIC into biomass via various carboxylation reactions involved in central metabolic functions such as carbon assimilation, anaplerosis and/or redox-balancing (9, 10).

Below the sunlit surface layer, the downward flux of particulate organic material is considered the main source of organic carbon (C) sustaining the dark ocean’s heterotrophic food web (11). However, current estimates point to a mismatch between organic matter consumption and supply in the dark ocean (12, 13), implying that additional sources of organic C are required to reconcile the C budget in the mesopelagic (defined here as the zone below the base of the euphotic zone) (14, 15). DIC fixation can be substantial in the dark ocean (14, 16) and often comparable in magnitude to microbial heterotrophic activities (14, 17, 18). In aphotic waters, DIC fixation has largely been attributed to the activities of chemolithoautotrophic bacteria and archaea that use the energy released from oxidizing reduced compounds to fuel a variety of inorganic carbon fixation pathways (14, 19–21). Chemolithoautotrophic production is a source of particulate organic C to deep waters that could contribute substantially to the microbial heterotrophic C demand in the dark ocean (14, 16). Additionally, chemolithoautotrophs release some of their recently fixed DIC as dissolved organic carbon (DOC) (22), likely further supporting microbial heterotrophic activities below the euphotic zone (23, 24). However, the activities of chemolithoautotrophic microbes are typically not well accounted for in mesopelagic C budgets (25, 26). An improved understanding of microbial processes in the mesopelagic is essential to better predict C export and sequestration, particularly under future climate change scenarios (27, 28).

Organic matter remineralization releases ammonium (29), which is the primary energy source for chemolithoautotrophy in most parts of the global ocean (19). Consequently, chemolithoautotrophic nitrification – the microbial oxidation of ammonia (NH_3_) to nitrite (NO_2-_) and further to nitrate (NO_3-_) – is expected to be tightly linked to dark DIC fixation rates below the euphotic zone. However, dark DIC fixation and nitrification are not routinely measured on oceanographic expeditions – particularly not in combination – making it impossible to accurately infer such a relationship. Empirical conversion factors obtained from pure culture studies are often used to estimate dark DIC fixation rates from nitrification rates (22, 30, 31) and *vice versa* (14, 32). Measurements of DIC fixation in the deep ocean are on average one order of magnitude higher than what could be supported by ammonium supplied by the sinking flux of particulate organic nitrogen (N), based on estimates of global ocean N export (19, 22, 30). To address this large discrepancy, multiple hypotheses have been proposed, including 1) unaccounted sources of ammonium to the deep ocean, 2) energy sources other than ammonium support chemolithoautotrophy, and 3)heterotrophic microbes are a major contributor to dark DIC fixation (15). However, direct evidence supporting any of these scenarios is thus far lacking.

In this study, we established a methodological framework to specifically inhibit ammonia-oxidizing microorganisms in ocean samples and confirmed that the inhibitor phenylacetylene had no measurable effects on other community-level processes. We then quantified the fraction of dark DIC fixation fueled by ammonia oxidation during two oceanographic expeditions in the eastern tropical and subtropical Pacific Ocean. Finally, we explored the relationships between rates of dark DIC fixation, nitrification and heterotrophic production. The results of this study help to reconcile the observed discrepancies between N supply and DIC fixation at depth, and advance our understanding of microbial processes in the mesopelagic ocean.

## Results and Discussion

### Environmental context

We sampled the eastern tropical and subtropical Pacific Ocean at eight stations spanning 35º N to 10º S during two oceanographic sampling campaigns (Fig. 1a). Sampling stations included regions of varying productivity, from productive coastal waters (Stn 1 and 3), the Equatorial upwelling region (Stn 10), to oligotrophic offshore stations in the Eastern Tropical North Pacific (ETNP; Stn 6, 12 and 15) and South Pacific (ETSP; Stn 5 and 7). A thick oxygen-deficient zone (ODZ; O_2_ 10 µM) was present at most stations, with the narrowest ODZ of ∼50 m found at the equatorial station (Stn 10) (*SI Appendix*, Fig. S1). We observed pronounced deep chlorophyll maxima (DCM) at all offshore stations (Fig. 1b) and secondary chlorophyll maxima at the two most northern offshore ETNP stations (Stn 6 and 15), reaching as deep as 125 m at Stn 15. Nitrification rates ranged between 0.4 and 64 nM d^-1^, showed typical maxima close to the base of the euphotic zone and sharply declined towards the mesopelagic, except at the most coastal station (Stn 1) where nitrification rates remained comparably high in deeper waters (Fig. 1b).

**Figure 1.**
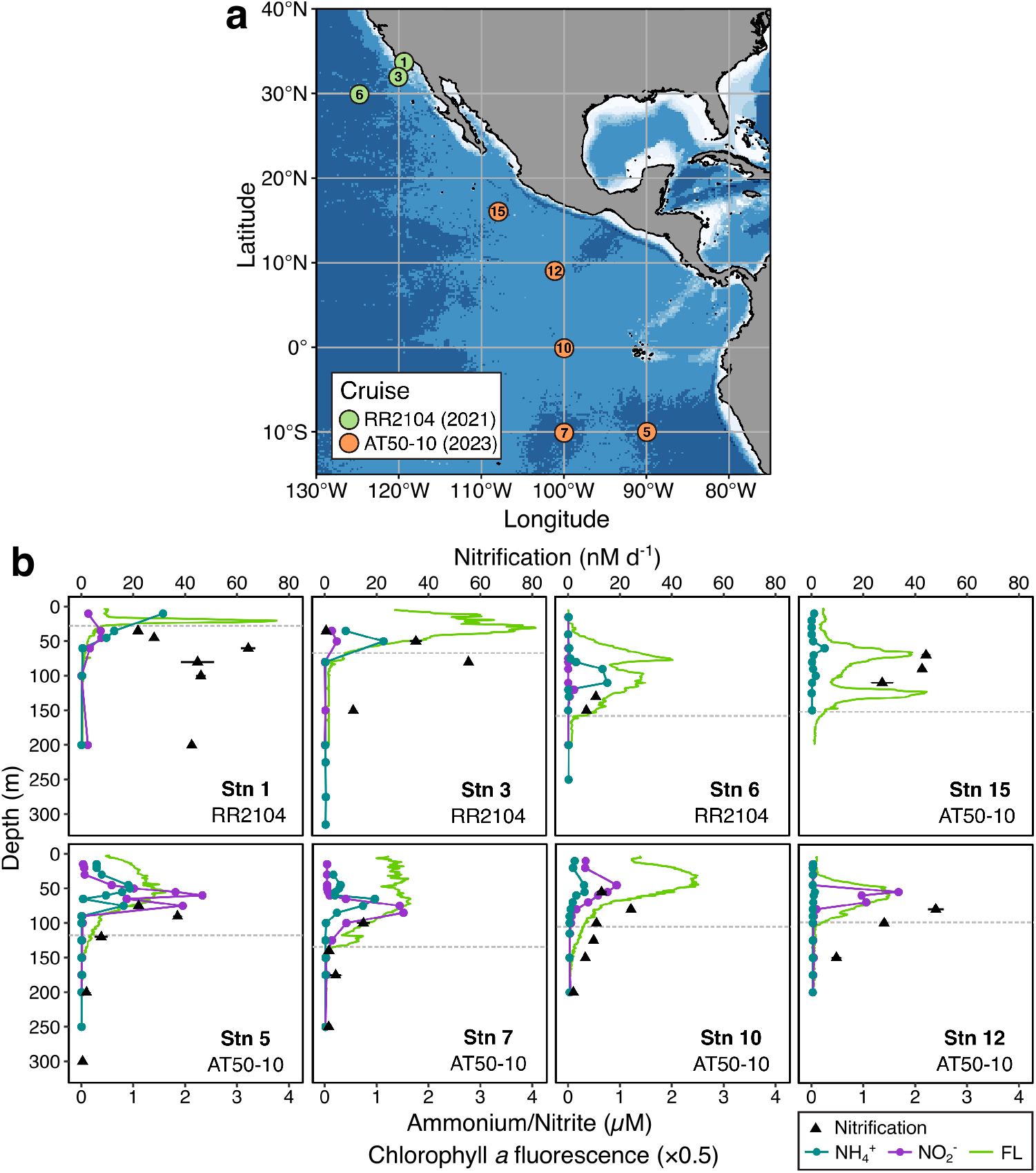
Cruise tracks and environmental context. **a)** Station map of cruises RR2104 (green) and AT50-10 (orange). **b)** Nitrification rates, chlorophyll *a* fluorescence (FL), ammonium (NH^4+^) and nitrite (NO_2-_) concentrations at eight stations during cruises AT50-10 and RR2104. NO_2-_ concentrations were not measured at Stn 15. For nitrification rates, the mean and standard deviation of triplicate measurements are shown. Depth of the euphotic zone is indicated by gray dashed horizontal lines.

AOA of the *Nitrosopumilaceae* were the main ammonia oxidizers at all stations and depths, making up 7-23% of the microbial community below the euphotic zone (Fig. 2), while bacterial ammonia oxidizers made up ≤ 0.15% of the microbial community throughout the water column. AOA abundances were low in surface waters and sharply increased to ∼10^7^ cells L^-1^ at the base of the euphotic zone, coinciding with high nitrification rates and low to undetectable (<10 nM) ammonium concentrations (Fig. 1b). Within the mesopelagic, AOA abundances remained relatively constant, except for anoxic depths (<1 µM O_2_) where their abundances declined substantially (*e.g*., Stn 12, 600 m depth). *Candidatus* Nitrosopelagicus (Water Column A Clade, WCA) was the dominant AOA genus in shallow waters, while members of the Water Column B (WCB) Clade dominated below 100 m depth and were often the only ammonia oxidizers in the mesopelagic (Fig. 2), in agreement with previous observations in the oligotrophic ocean (33). *Nitrosopumilus* 16S rRNA gene sequences were detected in shallow waters at Stn 5 and 15, but not within the mesopelagic. Estimated 16S rRNA gene abundances of *Ca*. Nitrosopelagicus were positively correlated with nitrification rates (*R*^2^=0.47, *SI Appendix*, Fig. S2a), as reported previously for WCA *amoA* genes and nitrification rates in the ETSP (*R*^2^=0.61, (34)). In contrast, WCB Clade abundances were negatively correlated with nitrification rates (*R*^2^=0.43, *SI Appendix*, Fig. S2b), suggesting that *Ca*. Nitrosopelagicus were the main contributors to ammonia oxidation at our study sites.

**Figure 2.**
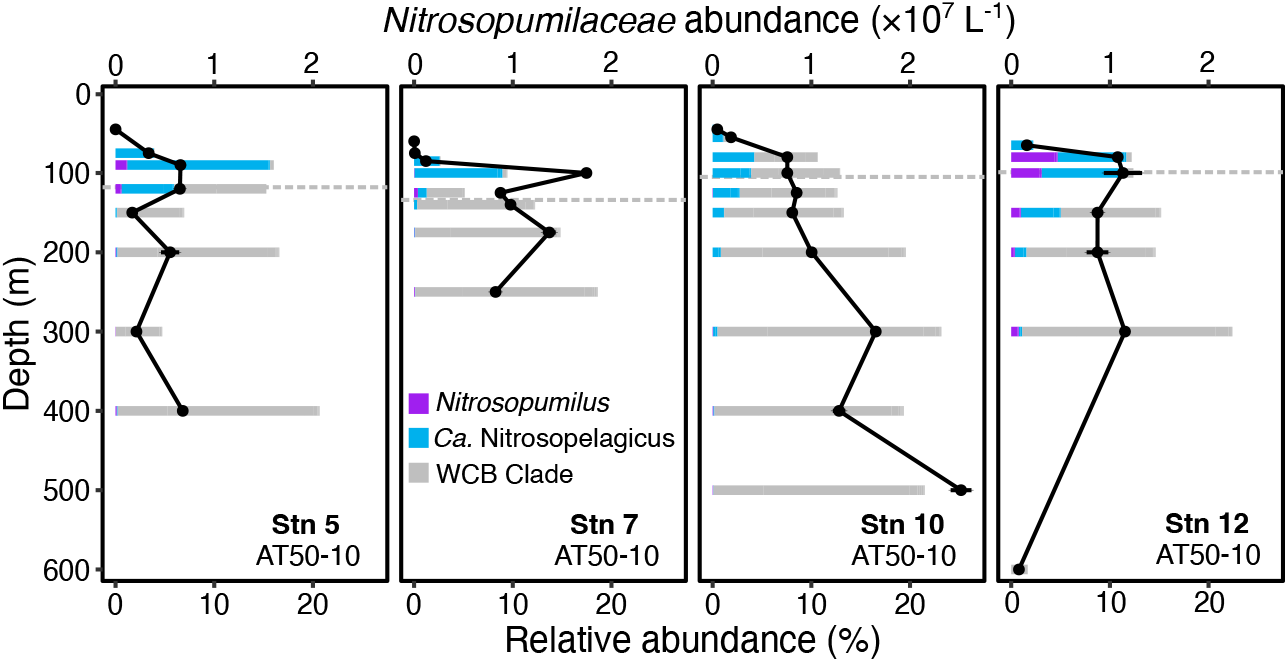
*Nitrosopumilaceae* abundances and community composition at four stations in the eastern Pacific (cruise AT50-10). Relative 16S rRNA gene abundances of different AOA clades are shown as a fraction of the total microbial community. Absolute *Nitrosopumilaceae* abundances were derived from quantitative PCR assays. Depth of the euphotic zone is indicated by gray dashed horizontal lines.

### Specific inhibition of ammonia oxidation by phenylacetylene

A reliable method to specifically inhibit the activities of ammonia oxidizers in ocean samples is required to isolate their contribution to dark DIC fixation. Phenylacetylene has previously been shown to inhibit cultures of terrestrial ammonia oxidizers by irreversibly binding to the ammonia monooxygenase enzyme (35), and is more practical for handling on oceanographic expeditions than the well-characterized inhibitor acetylene gas (36, 37). We first determined the effective inhibitory concentration of phenylacetylene on nitrification and DIC fixation activities of marine ammonia oxidizer cultures (≥5 µM, *SI Appendix*, Fig. S3). We further evaluated off-target effects of phenylacetylene on other members of the microbial community. Phenylacetylene is known to inhibit other monooxygenase enzymes, including soluble and particulate methane monooxygenases (38, 39). However, methane monooxygenases are only inhibited at 10 to 100 times higher effective concentrations (39) and relative abundances of putative methanotrophs were negligible at our study sites (<0.02%, Data File S1). Phenylacetylene must be dissolved in dimethylsulfoxide (DMSO) due to its low solubility in water, which could further affect microbial activities; we therefore tested the effect of DMSO separately on community-level process rates at oceanographic stations spanning contrasting productivity regimes and depth layers (Fig. 3). When 10 µM phenylacetylene in DMSO was added to whole seawater, nitrification rates were completely inhibited at all tested stations, while NO_2_^-^ oxidation and microbial heterotrophic production rates did not differ from those measured in control incubations without phenylacetylene (Fig. 3a and b). We further confirmed that the applied DMSO concentration alone had no measurable effect on microbial heterotrophic production and dark DIC fixation rates during the timeframe of our experiments (Fig. 3b and c). Consequently, phenylacetylene appears to be an effective, specific inhibitor of ammonia oxidation activity in the ocean and can be used to infer the contributions of ammonia oxidizers to dark DIC fixation rates (=“ammonia-fueled dark DIC fixation”).

**Figure 3.**
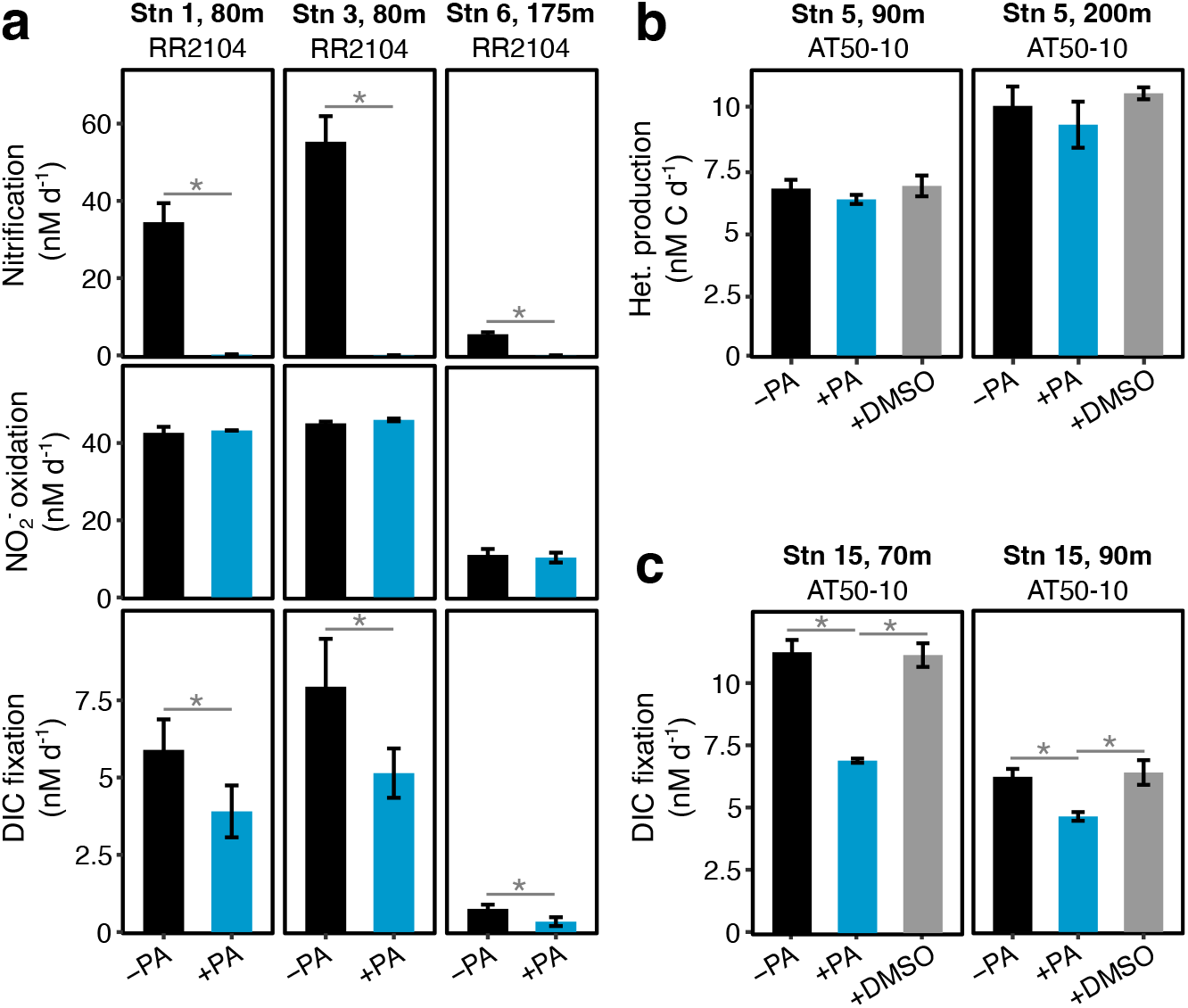
Effect of phenylacetylene (PA) on microbial process rates in environmental samples. **a**) Effect of 10 µM phenylacetylene additions on nitrification, nitrite oxidation and DIC fixation rates at stations 1, 3 and 6 during cruise RR2104. Comparison between additions of phenylacetylene (10 µM) dissolved in DMSO (0.01%) or DMSO alone on (**b**) microbial heterotrophic production, and (**c**) DIC fixation rates at stations 5 and 15 during cruise AT50-10, respectively. Significant differences between treatments (Student’s *t*-test, *P* < 0.05) are indicated by an asterisk (*). A Bonferroni correction was applied when multiple comparisons were made (panels b and c).

### Total and ammonia-fueled dark DIC fixation rates in the eastern tropical Pacific

We measured total dark DIC fixation rates throughout the water column of four stations, spanning the ETNP, ETSP and Equatorial Pacific (Fig. 4). Dark DIC fixation rates decreased with depth from the euphotic zone to the upper mesopelagic, with highest rates of ∼11 nM d^-1^ observed at 65 m (Stn 12) and 70 m (Stn 15) depth in the ETNP (Fig. 3c and 4). Throughout the mesopelagic, dark DIC fixation rates ranged between 0.3-1.8 nM d^-1^, with slightly increased rates in anoxic waters (O_2_ <1 µM) at Stn 5 and 12, possibly due to the activities of anaerobic chemolithoautotrophs (40, 41). Depth-integrated dark DIC fixation rates ranged between 0.2-0.8 mmol C m^-2^ d^-1^, which is considerably lower than in the North Atlantic (1.8 to 3.2 mmol C m^2^ d^1^, (14)). Compared to the eastern tropical Pacific, organic matter export from the euphotic zone is estimated to be higher in the North Atlantic (42), leading to higher overall productivity and possibly higher ammonium availability, which could explain these observed differences.

**Figure 4.**
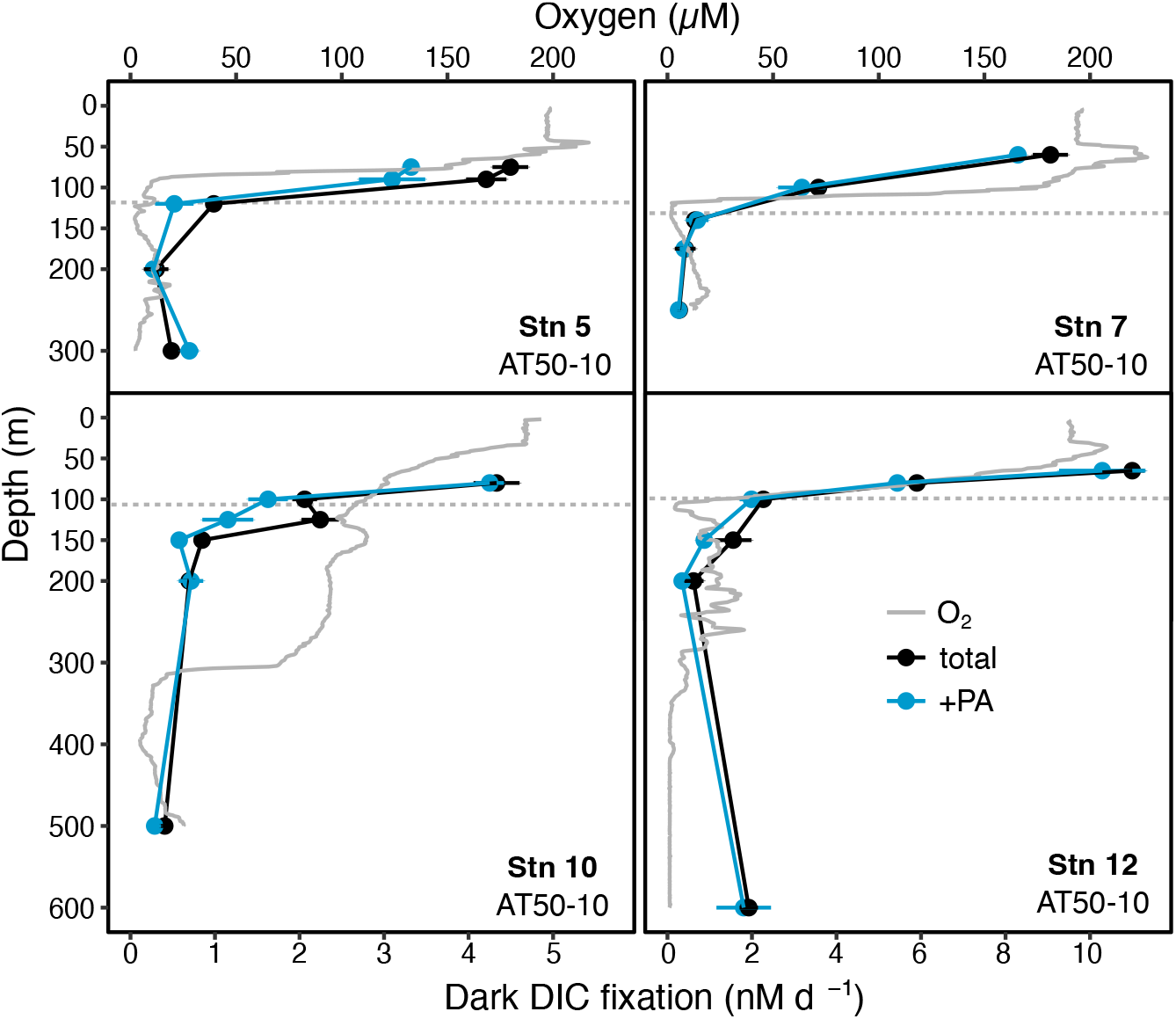
Depth-resolved dark DIC fixation rates at four stations in the eastern tropical Pacific during cruise AT50-10. Total dark DIC fixation rates are depicted in black and dark DIC fixation rates after phenylacetylene addition (+PA) in blue. Mean and standard deviations of triplicate measurements are shown. Oxygen concentration profiles are depicted in gray. Note that the scales on the x-axis are different between different stations.

We further determined the contribution of ammonia oxidizers to total dark DIC fixation in the epipelagic and upper mesopelagic. When phenylacetylene was added to incubation bottles, dark DIC fixation rates decreased on average by 33% (sd=18%, *n*=11) below the base of the euphotic zone (Fig. 4), with the exception of Stn 7, where surprisingly no decrease was observed. Within the euphotic zone, dark DIC fixation rates decreased on average by 18% (sd=12%, *n*=11), suggesting that ammonia oxidizers contribute less to dark DIC fixation within the euphotic zone compared to waters below. Ammonia oxidizers are inhibited by light (43–46) and euphotic zone nitrification rates are typically low under *in situ* light conditions (34, 47–49). However, AOA were abundant within the lower euphotic zone in our samples (Fig. 2), and incubations were performed in the dark, excluding the possibility of acute light inhibition. In deeper waters (>200 m), we only observed a decrease (29%) in dark DIC fixation at the Equator (Stn 10), while deep rates were not affected by phenylacetylene at the other three stations (Fig. 4). We hypothesize that this could possibly be due to the thick ODZ observed at all stations except for the equatorial Stn 10 (*SI Appendix*, Fig. S1), and the reliance of ammonia oxidation on O_2_ availability (50, 51).

Dark DIC fixation appeared to be slightly stimulated by phenylacetylene addition at Stn 5 at 300 m depth (Fig. 4). We cannot rule out the possibility that DMSO might serve as an alternative electron acceptor for anaerobic respiration after depletion of nitrate (52). However, stimulation of DIC fixation after phenylacetylene addition was not observed at other stations or depths, and nitrate concentrations remained high (≥30 µM) throughout the mesopelagic (Data File S2). Alternatively, phenylacetylene could potentially be used as an energy source supporting heterotrophic growth. However, since heterotrophic production was not stimulated by phenylacetylene addition (Fig. 3b), it is unlikely to be a commonly used substrate for microbes in the dark ocean.

Overall, ammonia oxidation could account for only 2 to 22% of the depth-integrated dark DIC fixation rates in the eastern tropical Pacific (Fig. 4). This implies that other microbial metabolisms contribute substantially to the cycling of inorganic carbon in the oceanic water column. Consequently, dark DIC fixation rates cannot be used to infer nitrification rates. In contrast, ammonia-fueled dark DIC fixation rates could be inferred from nitrification rates if suitable conversion factors were available.

### DIC fixation yields of ammonia oxidizers in the ocean

We aimed to better constrain DIC fixation yields (moles C fixed per mole of N oxidized) of ammonia oxidizers in the ocean to improve conversion factors for biogeochemical models. Nitrification and the fraction of dark DIC fixation inhibited by phenylacetylene (=“ammonia-fueled dark DIC fixation”) were positively correlated (*R*^2^=0.59, Fig. 5a). Average DIC fixation yield derived from the slope of the regression was ∼0.05, which is considerably lower than for cultures of *Ca*. Nitrosopelagicus brevis U25 and *Nitrosopumilus* sp. CCS1 isolated from the North Pacific (mean±sd: 0.09±0.01, (22)). The lower observed DIC fixation yields of AOA in the ocean might suggest a lower metabolic efficiency in the environment compared to ideal culture conditions.

**Figure 5.**
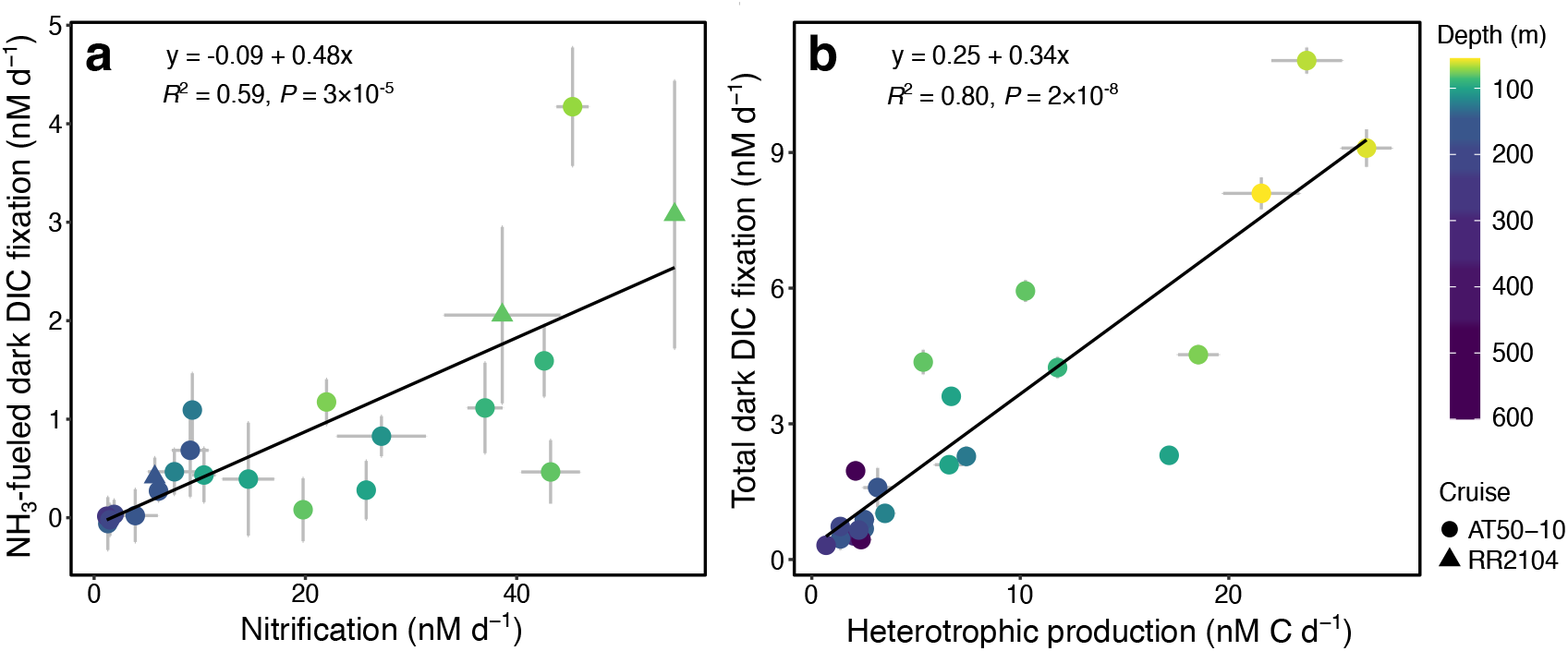
Relationships between dark DIC fixation, nitrification and heterotrophic production rates. **a)** Correlation between nitrification rates and the fraction of dark DIC fixation inhibited by phenylacetylene (=“ammonia-fueled dark DIC fixation”). NH_3_-fueled dark DIC fixation rates were calculated from subtracting triplicate phenylacetylene-inhibited measurements from triplicate non-inhibited measurements, error bars show the propagated error. **b)** Correlation between heterotrophic production and total dark DIC fixation rates. Error bars show the standard deviation of triplicate measurements. One data point (Stn 5, 200 m depth) was excluded from the plot due to unrealistically high heterotrophic production rates, possibly resulting from contamination or human error. Sample depth is color-coded with darker colors reflecting deeper depths. The two different cruises are differentiated by symbols (circle = AT50-10, triangle = RR2104). Heterotrophic production was only measured during cruise AT50-10.

Possible stimulation of nitrification due to ^15^N-ammonium tracer additions is a relevant concern for environmental rate measurements. While our tracer additions were relatively low (70-200 nM ^15^N-ammonium), we cannot rule out possible stimulation of ammonia oxidation, particularly in samples for which we measured high nitrification rates but did not observe any inhibition by phenylacetylene (i.e., Stn 10, 80 m depth; Stn 12, 80 and 100 m depth). A potential stimulation of ammonia oxidation may lead to lower observed DIC fixation yields when measuring rates from separate incubation bottles, with DIC bottles not receiving corresponding ammonium addition. This may have been the case on cruise AT50-10, where radioactivity safety protocols precluded combined rate measurements of nitrification and DIC fixation. In contrast, on cruise RR2104 ^13^C-labeled bicarbonate was used to measure DIC fixation rates in more productive waters, allowing us to combine both measurements within the same incubation bottle. Even when only considering samples from cruise RR2104, the average DIC fixation yields (0.054) were almost identical to those calculated from all data points (0.048). We therefore consider DIC fixation yields of ammonia oxidizers in this study realistic environmental estimates for the eastern tropical and subtropical Pacific, which can be used to inform theoretical models to better constrain the relationship between C and N fluxes in the dark ocean.

### Additional metabolisms contributing to dark DIC fixation

Ammonia is considered the primary energy source fueling water column chemolithoautotrophy (19), due to the higher molar ratio of N in marine organic matter relative to other potential energy sources such as reduced sulfur (S) and iron (53, 54). Ammonia oxidizers supply nitrite (NO_2_^-^) to nitrite oxidizers, and the two steps in the nitrification process are typically tightly coupled (55). Consequently, the inhibition of ammonia oxidation likely also results in the indirect inhibition of nitrite oxidation in the absence of an alternative NO_2_^-^ source. However, most of our stations were located in regions with pronounced ODZs, suggesting that NO_2_^-^ could also be supplied via nitrate reduction (56). We assessed nitrite oxidation rates at selected depths at four stations (*SI Appendix*, Fig. S4) and used average DIC fixation yields of cultured marine nitrite oxidizers (0.036, (22)) to estimate their potential contributions to dark DIC fixation. We estimate that ammonia and nitrite oxidizers together could account, on average, for 82% (sd=30%, *n*=11) of dark DIC fixation below the euphotic zone, but only 40% (sd=4%, *n*=5) within the deeper portion of the euphotic zone. In deeper waters (>200 m), the estimated contribution of both nitrifiers to dark DIC fixation might be more variable as suggested by the limited data available (21% at 300 m, Stn 5 and 63% at 250 m, Stn 7), which could also result from differences in the nitrite oxidizer community composition (*SI Appendix*, Fig. S5). However, given the lower DIC fixation yields observed for ammonia oxidation in the environment compared to those of cultured ammonia oxidizers (Fig. 5a, (22)), the contributions of nitrite oxidizers to dark DIC fixation could be substantially lower than estimated here.

The potential for sulfur oxidation-fueled chemolithoautotrophy is widespread in the dark ocean (20, 57, 58), and putative sulfur oxidizers were present at all stations (*SI Appendix*, Fig. S6). The organic S content of sinking particulate organic matter is ∼17-times lower than N (53), suggesting a very limited supply of reduced sulfur to the dark ocean. Even when considering the higher DIC fixation yields of sulfide oxidizers (0.15-0.35, (59–61)) compared to ammonia oxidizers (0.048, Fig. 5a), we estimate that sulfur-fueled chemolithoautotrophy could only amount to one third that of ammonia-fueled chemolithoautotrophy (*SI Appendix*, Table S1).

Heterotrophic microbes may also contribute to dark DIC fixation in the ocean (15, 18, 62), particularly in surface waters where uptake of DIC by different heterotrophic taxa has previously been shown to be high (40-200 nM C d^-1^), (63)). A compilation of >700 microbial heterotrophic production and dark DIC fixation rates from the Atlantic and Pacific Ocean indicated a positive correlation between both processes (*R*^2^=0.45, (15)). We found a similar, yet even stronger relationship between microbial heterotrophic production and total dark DIC fixation at four stations in the eastern tropical Pacific (*R*^2^=0.80, Fig. 5b). The few available data on heterotrophic DIC fixation suggest that 1–10% of carbon in bacterial biomass is derived from DIC assimilation (64–66). When assuming that heterotrophs incorporate 10% of their carbon as DIC, we could explain on average 30% (sd=15%, *n*=22) of dark DIC fixation rates in the epi- and upper mesopelagic at our study sites (*SI Appendix*, Table S1).

We estimate that nitrification, sulfur oxidation and heterotrophic DIC fixation together could account for 58-113% of the depth-integrated dark DIC fixation rates at our study sites, albeit the contributions of ammonia oxidizers are the first estimates derived from direct measurements.

## Conclusions

We established a methodological framework to robustly quantify the contribution of ammonia oxidizers to dark DIC fixation in the ocean. Our data confirm that ammonia oxidation is an important process in the epi- and upper mesopelagic, but we show that it contributes a much lower percentage of dark DIC fixation than would be predicted based on the stoichiometry of sinking organic matter flux, amounting to 2-22% of the depth-integrated rates in the eastern tropical and subtropical Pacific Ocean. When including high-end estimates of nitrite- and sulfur-fueled chemolithoautotrophy and heterotrophic DIC fixation, in total 58-113% of the depth-integrated dark DIC fixation rates could be explained (*SI Appendix*, Table S1). Discrepancies remain particularly within the euphotic zone (≤80 m depth), where contributions of ammonia and nitrite oxidizers to total dark DIC fixation are comparably small, and in deeper waters (≥200 m depth), where the flux of particulate organic matter from the surface is often insufficient to provide the required energy sources to sustain measured dark DIC fixation rates at depth. Constraining the contributions of sulfur oxidizers and heterotrophs will be crucial to reconcile these observed discrepancies.

Our results improve our understanding of microbial processes in the dark ocean and give new insights into the main energy sources fueling dark DIC fixation in the ocean. Our findings have broad implications by providing critical conversion factors for biogeochemical models of the mesopelagic carbon budget, which is essential to better predict the impact of future climate scenarios on the biological sequestration of carbon in the ocean.

## Materials and Methods

### Cruise track and dissolved nutrient analyses

Water samples were collected during two oceanographic cruises in the eastern tropical and subtropical Pacific Ocean aboard the R/V *Roger Revelle* (cruise RR2104: June 15^th^ – June 29^th^ 2021) and the R/V *Atlantis* (CliOMZ, cruise AT50-10: May 2^nd^ - June 9^th^ 2023) at a total of eight stations spanning 35º N to 10ºS (Fig. 1a). On both cruises, hydrographic data were collected with an SBE-9 plus conductivity, temperature, depth (CTD) sensor package (SeaBird Scientific) additionally equipped with a fluorometer (ECO, Seabird Scientific) and a Clark-type electrode oxygen sensor (SBE 43, SeaBird Scientific). Discrete water samples were collected using a rosette sampler equipped with 24 × 10 L Niskin bottles.

Ammonium (NH_4_^+^) concentrations were measured on board from unfiltered 40 mL seawater samples using the *o*-phthaldialdehyde derivatization method (67) with modifications as suggested in Taylor *et al*. (68) on an Aquafluor 8000 handheld fluorometer (Turner Designs). NH_4_^+^ standards (30 – 300 nM) were freshly prepared for each station using deep water (> 500 m), which consistently had a lower blank than ultrapure water. Samples for nitrite and nitrate concentration measurements were syringe-filtered (0.22 µm, Sterivex) and stored at –20°C before concentrations were determined by Cd reduction coupled to colorimetric detection via the Griess assay on a QuikChem 8500 Series 2 flow injection analysis system (Lachat Instruments) following established protocols (69). Euphotic zone depth was calculated as the depth below the chlorophyll *a* (Chl *a*) maximum, where Chl *a* fluorescence was 10% of the maximum fluorescence value (70).

### Nitrification rates

Nitrification and nitrite oxidation rates were determined using ^15^N isotope tracer methods. For depths where O_2_ concentrations were >20 µM, incubations were conducted in 250 mL or 1 L polycarbonate bottles (Nalgene). For depths where O_2_ concentrations were ≤ 20 µM, incubations were either conducted in 500 mL Tedlar bags (Restek) or 120 mL glass serum bottles following procedures to prevent O_2_ contamination (*SI Appendix*). For each incubation depth, three bottles or bags were filled and spiked with [^15^N]-tracer (99 atm% ^15^NH_4_Cl or 98 atm% Na^15^NO_2_^-^, Cambridge Isotope Laboratories) to a final label concentration of 70 to 200 nM, depending on the depth and productivity region the sample was collected from.

Nitrification rates were determined throughout the water column targeting depths in the middle of the euphotic zone, the nitrification maximum and the upper mesopelagic. Nitrite oxidation rates were measured during cruise AT50-10 at a lower resolution compared to nitrification rates (*SI Appendix*, Fig. S4). Incubations were conducted within ±1.5°C of *in situ* temperature. At timepoints of 0, 12, and 24 h, 50 mL samples were drawn from each bottle or bag, 0.2 µm syringe-filtered into a 20 mL HDPE bottle, and frozen at -20ºC. Frozen samples were transported to the laboratory, thawed, and prepared for *δ*^15^N_NO2+NO3_ analysis using the ‘denitrifier method’ (71). For nitrite oxidation rate samples, the added ^15^NO_2_^-^ tracer was removed using sulfamic acid addition and subsequent neutralization with NaOH prior to sample preparation (72). Samples were analyzed on a custom purge and trap system interfaced to a Thermo Delta Plus XP isotope ratio mass spectrometer (73). *δ*^15^N_NOx_ values were calibrated against NO_3_^-^ isotope reference materials USGS 32, USGS 34, and USGS 35, analyzed in parallel. Nitrification and nitrite oxidation rates were calculated using the basic equations of Dugdale and Goering (74).

### Dark DIC fixation rates

During cruise RR2104, DIC fixation was measured via the incorporation of [^13^C]-bicarbonate. Water was sampled from Niskin bottles into 1 L polycarbonate bottles (Nalgene). For each station, the depth of the expected nitrification maximum was sampled, and twelve bottles were filled to which [^13^C]-bicarbonate and either [^15^N]-ammonium or [^15^N]-nitrite were added (see section on nitrification rates). [^13^C]-bicarbonate tracer concentrations were made as 20% additions of the ambient bicarbonate pool. Phenylacetylene (10 µM) dissolved in dimethylsulfoxide (DMSO, 0.01%) was added to six of the incubation bottles to inhibit ammonia oxidation activities. After 24 h of incubation, samples were filtered onto pre-combusted glass fiber filters (GF75, 25mm, Advantec) and frozen at -80°C until further processing. Filters were acidified by acid fumigation to remove inorganic carbonates and dried at 60°C for at least 24 h. δ^13^C of particulate organic carbon (POC) was measured with a Finnigan Delta-Plus Advantage isotope ratio mass spectrometer (Thermo Scientific) coupled with an elemental analyzer (Costech EAS). Acetanilide reference standards were run at the beginning of each set of 35 samples and tested every 5 samples within each set. Instrument precision, determined using replicate analyses of L-glutamic acid USGS 40, was ±0.1‰ for ^13^C. δ^13^C of DIC in water samples was measured with a GasBench II system interfaced to a Delta V Plus isotope ratio mass spectrometer (Thermo Scientific) at the UC Davis Stable Isotope Facility. DIC fixation rates were calculated as described previously (75).

During cruise AT50-10, DIC fixation was measured via the incorporation of [^14^C]-bicarbonate as previously described (76), with modifications. For depths where O_2_ concentrations were >20 µM, water was sampled into 1 L polycarbonate bottles (Nalgene) and later dispersed into 40 mL glass vials with teflon coated silicon septa (TOC-certified, Fisher Scientific). Initially, we used 50 mL conical centrifuge tubes (Fisher Scientific) for oxic incubations, however, we noticed that DIC fixation rates were highly variable and much higher in plastic compared to glass tubes (*SI Appendix*, Fig. S7). Consequently, none of the data obtained from measurements in plastic tubes could be used for further analysis, and we switched to glass tubes by Stn 5. For depths where *in situ* O_2_ concentrations were ≤ 20 µM, water was sampled directly from the Niskin bottle into 60 mL glass serum bottles following procedures to avoid O_2_ contamination (*SI Appendix*).

For each incubation depth, seven or eight replicate bottles were filled and spiked with 50 µCi [^14^C]-bicarbonate (specific activity 56 mCi mmol^−1^ / 2.072 × 10^9^ Bq mmol^−1^; PerkinElmer). To three bottles, 10 µM phenylacetylene dissolved in DMSO (0.01% final concentration) was added to inhibit ammonia oxidation activities. One to two bottles served as killed controls to which formaldehyde (3% vol/vol) was added at the start of incubation. Bottles were incubated in the dark at *in situ* temperature and handled under red light to prevent ^14^C assimilation by phytoplankton. Live incubations were terminated after 24-72 h by adding formaldehyde (3% vol/vol), and after 30–60 min, samples were filtered onto 0.2 *μ*m polycarbonate filters (GTTP, 25 mm, Millipore) and rinsed with 10 mL of artificial seawater. The filters were transferred to scintillation vials and 10 mL of scintillation cocktail (Ultima Gold; PerkinElmer) was added. Samples were shaken for ca. 30 s and placed in the dark for at least 24 h prior to counting the disintegrations per minute (DPM) in a scintillation counter (Perkin-Elmer Tri-Carb 2910 TR) for 15 min. Total radioactivity measurements were performed to verify added [^14^C]-bicarbonate concentrations by pipetting 100 *μ*L of unfiltered sample into scintillation vials containing 400 *μ*L beta-phenylethylamine, and samples were immediately measured after addition of scintillation cocktail. DIC fixation rates were calculated as described previously (22).

Due to the different methodologies used on both cruises to infer DIC fixation rates, we compared rates obtained from [^13^C]- and [^14^C]-bicarbonate incorporation on a culture of the ammonia-oxidizing archaeon *N. adriaticus* NF5 and found a good agreement between the two methods (*SI Appendix*, Fig. S8a). We further confirmed that [^14^C]-bicarbonate incorporation rates were linear over the length of the incubation time (up to 3 days, *SI Appendix*, Fig. S8b).

### Leucine incorporation rates

Microbial heterotrophic production rates were measured via the incorporation of [^3^H]-leucine into microbial biomass using a modified version of the microcentrifuge method (77). Briefly, [^3^H]-leucine (specific activity 44.9 Ci mmol^-1^; Perkin Elmer) was added to 1.6 mL of sample at a final concentration of 20 nM and incubated for 2-3 h at *in situ* temperature. For each depth and station, triplicate live incubations and one killed control, to which 100 µL of trichloroacetic acid (TCA) was added immediately, were carried out. Incubations were terminated by addition of cold 100 µL 100% TCA and stored at 4°C until extraction. Proteins were extracted following the procedures described in (78), 1.5 mL scintillation cocktail (Ultima Gold, Perkin Elmer) was added to each tube, and DPM were measured on a scintillation counter (Perkin-Elmer Tri-Carb 2910 TR) for 2 min. [^3^H]-leucine incorporation rates were converted to units of carbon using a conversion factor of 1.5 kg C (mol leucine incorporated)^-1^ (79). To test the effects of phenylacetylene (10 µM) and DMSO (0.01%) on microbial heterotrophic production, [^3^H]-leucine was added to a final concentration of 60 nM and incubated for 24h. All other steps followed the procedures described above.

### Quantitative PCR and metagenome analysis

Seawater was collected into acid-washed 4 L polycarbonate bottles, biomass was sequentially filtered onto 5 µm (25 mm, PETE, Sterlitech Corporation) and 0.22 µm (25mm, Supor PES, Pall) pore size membrane filters, which were subsequently frozen at -80ºC until extraction. DNA was extracted from 0.22 µm filters using the Qiagen DNeasy Blood & Tissue kit with modifications as previously described (80).

Quantitative PCR assays were conducted using group-specific assays for the 16S ribosomal RNA (rRNA) gene of marine AOA and NOB of the *Nitrospinaceae* family following established protocols and thermocycling conditions (80, 81). We modified the previously published AOA primers (81) to increase the coverage of members of the *Nitrosopumilus* genus (MGI_F: 5’-GTC TAC CAG AAC ACG TYC-3’, MGI_R: 5’-WGG CGT TGA CTC CAA TTG-3’). Gene copies were quantified on a CFX96 qPCR machine (Bio-Rad) with SYBR Green chemistry. All samples were run in triplicate against a standard curve spanning approximately 10^1^ – 10^6^ template copies. Linearized plasmids containing cloned inserts of the target genes (TOPO pCR4 vector, Invitrogen) were used as standards as described in (82). Fresh standard dilutions were made from frozen stocks for each day of analysis. Efficiencies of the assays were 99.5-100.5%.

DNA libraries were prepared by the UC Davis Genome Center and sequenced on two flowcells of the AVITI sequencer (Element Biosciences) run with paired-end 300 bp reads (detailed methodological descriptions in the *SI Appendix*). Resulting reads were trimmed, quality-filtered, and internal standards removed (83, 84). Microbial community composition was subsequently assessed by mapping reads to the SILVA SSU rRNA reference database (v138.2) using phyloFLASH (v 3.4.2) (85).

## Data Availability

Hydrographic data of cruises RR2104 and AT50-10 are available on the “Rolling deck to Repository” (R2R) webpage (https://www.rvdata.us) under the respective cruise names. Raw metagenome sequencing data generated during this study are available in the NCBI sequence read archive (SRA) repository under accession numbers SRR31156201-SRR31156268 and SRR31341463-SRR31341480 (BioProject: PRJNA1179712). PhyloFLASH analyses are available in Data File S1. Environmental nutrient and rate measurement data is available Data File S2 and on the BCO-DMO repository (https://www.bco-dmo.org) under project number 806565. All other data are available in the manuscript or *SI Appendix*.

## Acknowledgements

We thank Anela Choy, chief scientist of expedition RR2104, as well as the captain and crew of the R/V *Atlantis* and R/V *Roger Revelle*. We thank Christie Yorke for assistance with nutrient and nitrification rate measurements, and Ken Marchus for support with running the IRMS. We are grateful to Elisa Halewood and Thomas Reinthaler for their support and helpful discussions on procedures for ship-based radioisotope experiments, and Laura Bristow for advice on experimental setup and sharing equipment for low oxygen incubations. We would also like to thank Oliver Fajardo and his team at UC San Diego for their support in radioactive waste management. This research was supported by the Austrian Science Fund (FWF) Project J4426-B to BB and the US National Science Foundation (NSF) award OCE-1924512 and Simons Foundation award LI-SIAME-00001560 to AES. CAC was supported by the Simons Foundation International’s BIOS-SCOPE program. BB and KK were additionally supported by the FWF Wittgenstein Award Z-383B to MW. We acknowledge funding from the UC Ship Funds Program for cruise RR2104 (to Anela Choy). MW and KK also acknowledge funding from the FWF Cluster of Excellence “Microbiomes drive planetary health” (10.55776/COE7), and BB acknowledges funding from the European Union (ERC, METHANIAQ, 101116021). Views and opinions expressed are however those of the author(s) only and do not necessarily reflect those of the European Union or the European Research Council Executive Agency.

## References

1. C. B. Field, M. J. Behrenfeld, J. T. Randerson, P. Falkowski, Primary Production of the Biosphere: Integrating Terrestrial and Oceanic Components. Science 281, 237–240 (1998).

2. A. Prakash, R. W. Sheldon, W. H. Sutcliffe, Geographic variation of oceanic ^14^C dark uptake. Limnology & Oceanography 36, 30–39 (1991).

3. W. K. W. Li, B. D. Irwin, P. M. Dickie, Dark fixation of ^14^C: Variations related to biomass and productivity of phytoplankton and bacteria. Limnology and Oceanography 38, 483–494 (1993).

4. A. Alothman, D. López-Sandoval, C. M. Duarte, S. Agustí, Bacterioplankton dark CO_2_ fixation in oligotrophic waters. Biogeosciences 20, 3613–3624 (2023).

5. F. Baltar, G. J. Herndl, Ideas and perspectives: Is dark carbon fixation relevant for oceanic primary production estimates? Biogeosciences 16, 3793–3799 (2019).

6. H. Saxena, et al., Contribution of Carbon Fixation Toward Carbon Sink in the Ocean Twilight Zone.Geophysical Research Letters 49, e2022GL099044 (2022).

7. W. K. W. Li, P. M. Dickie, Light and dark ^14C^ uptake in dimly-lit oligotrophic waters: relation to bacterial activity. Journal of Plankton Research 13, 29–44 (1991).

8. L. Alonso-Saez, et al., Role for urea in nitrification by polar marine Archaea. Proc. Natl. Acad. Sci. U.S.A. 109, 17989–17994 (2012).

9. T. J. Erb, Carboxylases in natural and synthetic microbial pathways. Applied and Environmental Microbiology 77, 8466–8477 (2011).

10. A. Braun, et al., Reviews and syntheses: Heterotrophic fixation of inorganic carbon -Significant but invisible flux in environmental carbon cycling. Biogeosciences 18, 3689–3700 (2021).

11. T. Nagata, “Production mechanisms of dissolved organic matter” in Microbial Ecology of the Oceans, Wiley series in ecological and applied microbiology., D. L. Kirchman, Ed. (Wiley-Liss, 2000), pp. 121–152.

12. A. B. Burd, et al., Assessing the apparent imbalance between geochemical and biochemical indicators of meso- and bathypelagic biological activity: What the @$♯! is wrong with present calculations of carbon budgets? Deep Sea Research Part II: Topical Studies in Oceanography 57, 1557–1571 (2010).

13. G. J. Herndl, T. Reinthaler, Microbial control of the dark end of the biological pump. Nature Geoscience 6, 718–724 (2013).

14. T. Reinthaler, H. M. van Aken, G. J. Herndl, Major contribution of autotrophy to microbial carbon cycling in the deep North Atlantic’s interior. Deep-Sea Research Part II 57, 1572–1580 (2010).

15. G. J. Herndl, B. Bayer, F. Baltar, T. Reinthaler, Prokaryotic Life in the Deep Ocean’s Water Column.Annual Review of Marine Science 15, 461–483 (2023).

16. F. Baltar, et al., Significance of non-sinking particulate organic carbon and dark CO_2_ fixation to heterotrophic carbon demand in the mesopelagic northeast Atlantic. Geophysical Research Letters 37 (2010).

17. W. Zhou, et al., High dark carbon fixation in the tropical South China Sea. Continental Shelf Research 146, 82–88 (2017).

18. E. Guerrero-Feijóo, E. Sintes, G. J. Herndl, M. M. Varela, High dark inorganic carbon fixation rates by specific microbial groups in the Atlantic off the Galician coast (NW Iberian margin). Environmental Microbiology 20, 602–611 (2018).

19. J. J. Middelburg, Chemoautotrophy in the ocean. Geophysical Research Letters 38, 94–97 (2011).

20. B. K. Swan, et al., Potential for chemolithoautotrophy among ubiquitous bacteria lineages in the dark ocean. Science 333, 1296–1300 (2011).

21. M. G. Pachiadaki, et al., Major role of nitrite-oxidizing bacteria in dark ocean carbon fixation. Science 358, 1046–1051 (2017).

22. B. Bayer, K. McBeain, C. A. Carlson, A. E. Santoro, Carbon content, carbon fixation yield and dissolved organic carbon release from diverse marine nitrifiers. Limnology and Oceanography 68, 84–96 (2023).

23. B. Bayer, et al., Ammonia-oxidizing archaea release a suite of organic compounds potentially fueling prokaryotic heterotrophy in the ocean. Environmental Microbiology 21, 4062–4075 (2019).

24. B. Bayer, et al., Metabolite release by nitrifiers facilitates metabolic interactions in the ocean. The ISME Journal 18, wrae172 (2024). 10.1093/ismejo/wrae172.

25. S. L. C. Giering, et al., Reconciliation of the carbon budget in the ocean’s twilight zone. Nature 507, 480–483 (2014).

26. C. Baumas, et al., Reconstructing the ocean’s mesopelagic zone carbon budget: sensitivity and estimation of parameters associated with prokaryotic remineralization. Biogeosciences 20, 4165– 4182 (2023).

27. E. Y. Kwon, F. Primeau, J. L. Sarmiento, The impact of remineralization depth on the air-sea carbon balance. Nature Geoscience 2, 630–635 (2009).

28. A. J. Fassbender, et al., Amplified Subsurface Signals of Ocean Acidification. Global Biogeochemical Cycles 37, e2023GB007843 (2023).

29. D. L. Kirchman, “Introduction and Overview” in Microbial Ecology of the Oceans, (2008), pp. 1–26.

30. Y. Zhang, et al., Nitrifier adaptation to low energy flux controls inventory of reduced nitrogen in the dark ocean. Proc. Natl. Acad. Sci. U.S.A. 117, 4823–4830 (2020).

31. E. J. Zakem, et al., Controls on the relative abundances and rates of nitrifying microorganisms in the ocean. Biogeosciences 19, 5401–5418 (2022).

32. J. E. Dore, D. M. Karl, Nitrification in the euphotic zone as a source for nitrite, nitrate, and nitrous oxide at Station ALOHA. Limnology and Oceanography 41, 1619–1628 (1996).

33. A. E. Santoro, R. A. Richter, C. L. Dupont, Planktonic Marine Archaea. Annual Review of Marine Science 11, 131–158 (2019).

34. A. E. Santoro, et al., Nitrification and nitrous oxide production in the offshore waters of the Eastern Tropical South Pacific. Global Biogeochemical Cycles 35, 0e2020GB006716–3 (2021).

35. C. L. Wright, A. Schatteman, A. T. Crombie, J. C. Murrell, L. E. Lehtovirta-Morley, Inhibition of Ammonia Monooxygenase from Ammonia-Oxidizing Archaea by Linear and Aromatic Alkynes. Applied and Environmental Microbiology 86, e02388–19 (2020).

36. M. R. Hyman, D. J. Arp, 14C2H2- and 14CO2-labeling studies of the de novo synthesis of polypeptides by Nitrosomonas europaea during recovery from acetylene and light inactivation of ammonia monooxygenase. Journal of Biological Chemistry 267, 1534–1545 (1992).

37. M. R. Hyman, P. M. Wood, Suicidal inactivation and labelling of ammonia mono-oxygenase by acetylene. Biochemical Journal 227, 719–725 (1985).

38. W. K. Keener, et al., Use of selective inhibitors and chromogenic substrates to differentiate bacteria based on toluene oxygenase activity. Journal of Microbiological Methods 46, 171–185 (2001).

39. S. Lontoh, et al., Differential inhibition in vivo of ammonia monooxygenase, soluble methane monooxygenase and membrane-associated methane monooxygenase by phenylacetylene. Environmental Microbiology 2, 485–494 (2000).

40. T. Dalsgaard, B. Thamdrup, L. Farías, N. P. Revsbech, Anammox and denitrification in the oxygen minimum zone of the eastern South Pacific. Limnology and Oceanography 57, 1331–1346 (2012).

41. C. M. Callbeck, et al., Oxygen minimum zone cryptic sulfur cycling sustained by offshore transport of key sulfur oxidizing bacteria. Nature Communications 9, 1729 (2018).

42. M. Nowicki, T. DeVries, D. A. Siegel, Quantifying the Carbon Export and Sequestration Pathways of the Ocean’s Biological Carbon Pump. Global Biogeochemical Cycles 36, e2021GB007083 (2022).

43. A. B. Hooper, K. R. Terry, Photoinactivation of Ammonia Oxidation in Nitrosomonas. J Bacteriol 119, 899–906 (1974).

44. S. G. Horrigan, A. F. Carlucci, P. M. Williams, Light inhibition of nitrification in sea-surface films.Journal of Marine Research 39, 557–565 (1981).

45. S. N. Merbt, et al., Differential photoinhibition of bacterial and archaeal ammonia oxidation. FEMS Microbiology Letters 327, 41–46 (2012).

46. W. Qin, et al., Marine ammonia-oxidizing archaeal isolates display obligate mixotrophy and wide ecotypic variation. Proc. Natl. Acad. Sci. U.S.A. 111, 12504–12509 (2014).

47. B. B. Ward, R. J. Olson, M. J. Perry, Microbial nitrification rates in the primary nitrite maximum off southern California. Deep Sea Research Part A. Oceanographic Research Papers 29, 247–255 (1982).

48. R. E. A. Horak, et al., Relative impacts of light, temperature, and reactive oxygen on thaumarchaeal ammonia oxidation in the North Pacific Ocean. Limnology and Oceanography 63, 741–757 (2018).

49. J. M. Smith, F. P. Chavez, C. A. Francis, Ammonium uptake by phytoplankton regulates nitrification in the sunlit ocean. PLoS ONE 9, e108173 (2014).

50. W. Martens-Habbena, D. A. Stahl, Nitrogen metabolism and kinetics of ammonia-oxidizing archaea. Methods Enzymol 496, 465–487 (2011).

51. J. A. Kozlowski, M. Stieglmeier, C. Schleper, M. G. Klotz, L. Y. Stein, Pathways and key intermediates required for obligate aerobic ammonia-dependent chemolithotrophy in bacteria and Thaumarchaeota. The ISME Journal 10, 1836–1845 (2016).

52. D. A. Tebbe, et al., Microbial drivers of DMSO reduction and DMS-dependent methanogenesis in saltmarsh sediments. The ISME Journal 17, 2340–2351 (2023).

53. P. A. Matrai, R. W. Eppley, Particulate organic sulfur in the waters of the Southern California Bight.Global Biogeochemical Cycles 3, 89–103 (1989).

54. B. S. Twining, S. B. Baines, The Trace Metal Composition of Marine Phytoplankton. Annual Review of Marine Science 5, 191–215 (2013).

55. B. B. Ward, “Nitrification in the Ocean” in Nitrification, B. B. Ward, D. J. Arp, M. G. Klotz, Eds. (ASM Press, 2011), pp. 325–346.

56. P. Lam, M. M. M. Kuypers, Microbial nitrogen cycling processes in oxygen minimum zones. Annual Review of Marine Science 3, 317–345 (2011).

57. F. Baltar, et al., A ubiquitous gammaproteobacterial clade dominates expression of sulfur oxidation genes across the mesopelagic ocean. Nature Microbiology 8, 1137–1148 (2023).

58. A. L. Jaffe, R. S. R. Salcedo, A. E. Dekas, Abundant and metabolically flexible lineages within the SAR324 and gammaproteobacteria dominate the potential for rubisco-mediated carbon fixation in the dark ocean. Biorxiv. Available at: http://biorxiv.org/lookup/doi/10.1101/2024.05.09.593449

59. D. C. Nelson, B. B. Jørgensen, N. P. Revsbech, Growth Pattern and Yield of a Chemoautotrophic Beggiatoa sp. in Oxygen-Sulfide Microgradients. Appl Environ Microbiol 52, 225–233 (1986).

60. J. M. Klatt, L. Polerecky, Assessment of the stoichiometry and efficiency of CO_2_ fixation coupled to reduced sulfur oxidation. Front. Microbiol. 6 (2015).

61. D. Vasquez-Cardenas, F. J. R. Meysman, H. T. S. Boschker, A Cross-System Comparison of Dark Carbon Fixation in Coastal Sediments. Global Biogeochemical Cycles 34, e2019GB006298 (2020).

62. S. DeLorenzo, et al., Ubiquitous Dissolved Inorganic Carbon Assimilation by Marine Bacteria in the Pacific Northwest Coastal Ocean as Determined by Stable Isotope Probing. PLoS ONE 7, e46695 (2012).

63. L. Alonso-Sáez, P. E. Galand, E. O. Casamayor, C. Pedrós-Alió, S. Bertilsson, High bicarbonate assimilation in the dark by Arctic bacteria. The ISME Journal 4, 1581–1590 (2010).

64. J. I. Sorokin, On the carbon dioxide uptake during cell synthesis by microorganisms. Zeitschrift für Allg. Mikrobiologie 6, 69–73 (1966).

65. P. Roslev, M. B. Larsen, D. Jørgensen, M. Hesselsoe, Use of heterotrophic CO2 assimilation as a measure of metabolic activity in planktonic and sessile bacteria. Journal of Microbiological Methods 59, 381–393 (2004).

66. M. Spona-Friedl, et al., Substrate-dependent CO2 fixation in heterotrophic bacteria revealed by stable isotope labelling. FEMS Microbiology Ecology 96, fiaa080 (2020).

67. R. Holmes, A. Aminot, R. Kérouel, B. Hooker, B. Peterson, A simple and precise method for measuring ammonium in marine and freshwater ecosystems. Canadian Journal of Fisheries and Aquatic Sciences 56, 1801–1808 (1999).

68. B. W. Taylor, et al., Improving the fluorometric ammonium method: Matrix effects, background fluorescence, and standard additions. Journal of the North American Benthological Society 26, 167– 177 (2007).

69. B. Hales, A. Van Geen, T. Takahashi, High-frequency measurement of seawater chemistry: Flow-injection analysis of macronutrients. Limnol. Oceanogr. Methods 2, 91–101 (2004).

70. S. A. Owens, S. Pike, K. O. Buesseler, Thorium-234 as a tracer of particle dynamics and upper ocean export in the Atlantic Ocean. Deep Sea Research Part II: Topical Studies in Oceanography 116, 42– 59 (2015).

71. D. M. Sigman, et al., A bacterial method for the nitrogen isotopic analysis of nitrate in seawater and freshwater. Analytical Chemistry 73, 4145–4153 (2001).

72. J. Granger, D. M. Sigman, M. G. Prokopenko, M. F. Lehmann, P. D. Tortell, A method for nitrite removal in nitrate N and O isotope analyses: Nitrite removal. Limnol. Oceanogr. Methods 4, 205–212 (2006).

73. M. R. McIlvin, K. L. Casciotti, Technical updates to the bacterial method for nitrate isotopic analyses. Analytical Chemistry 83, 1850–1856 (2011).

74. R. C. Dugdale, J. J. Goering, Uptake of new and regenerated forms of nitrogen in primary productivity. Limnology and Oceanography 12, 196–206 (1967).

75. J. Grosse, P. Van Breugel, H. T. S. Boschker, Tracing carbon fixation in phytoplankton—compound specific and total ^13^C incorporation rates. Limnol. Oceanogr. Methods 13, 288–302 (2015).

76. G. J. Herndl, et al., Contribution of Archaea to total prokaryotic production in the deep Atlantic Ocean. Applied and environmental microbiology 71, 2303–2309 (2005).

77. D. C. Smith, F. Azam, A simple, economical method for measuring bacterial protein synthesis rates in seawater using. Marine microbial food webs 6, 107–114 (1992).

78. N. Baetge, et al., Bacterioplankton response to physical stratification following deep convection. Elementa: Science of the Anthropocene 10, 00078 (2022).

79. M. Simon, F. Azam, Protein content and protein synthesis rates of planktonic marine bacteria. Marine Ecology Progress Series 51, 201–213 (1989).

80. A. E. Santoro, K. L. Casciotti, C. A. Francis, Activity, abundance and diversity of nitrifying archaea in the central California Current. Environmental Microbiology 12, 1989–2006 (2010).

81. T. J. Mincer, et al., Quantitative distribution of presumptive archaeal and bacterial nitrifiers in Monterey Bay and the North Pacific Subtropical Gyre. Environmental Microbiology 9, 1162–1175 (2007).

82. A. E. Santoro, et al., Thaumarchaeal ecotype distributions across the equatorial Pacific Ocean and their potential roles in nitrification and sinking flux attenuation. Limnology and Oceanography 62, 1984–2003 (2017).

83. B. Langmead, S. L. Salzberg, Fast gapped-read alignment with Bowtie 2. Nature Methods 9, 357– 359 (2012).

84. A. M. Bolger, M. Lohse, B. Usadel, Trimmomatic: a flexible trimmer for Illumina sequence data.Bioinformatics 30, 2114–2120 (2014).

85. H. R. Gruber-Vodicka, B. K. B. Seah, E. Pruesse, phyloFlash: Rapid Small-Subunit rRNA Profiling and Targeted Assembly from Metagenomes. mSystems 5, 10.1128/msystems.00920-20 (2020).

