## Supplementary Information for "Contribution of ammonia oxidizers to inorganic carbon fixation in the dark ocean"

**This PDF file includes:**

Supplementary Material and Methods

Supplementary Results and Discussion (Figures S1 to S8, Tables S1 and S2)

Supplementary References

### Supplementary Material and Methods

#### *Procedures for setting up incubations under low $O_2$ conditions*

At depths where *in situ* oxygen ($O_2$) concentrations were ≤20 µM, we adhered to the following procedures to avoid $O_2$ contamination during sampling and to replicate *in situ* conditions as well as possible.

For DIC fixation rate incubations, water was sampled directly from the Niskin bottle into 60 mL glass serum bottles using tygon tubing, allowing approximately three volumes of sample water to overflow the bottle prior to collection. Serum bottles were closed bubble-free with deoxygenated butyl rubber stoppers (1) and sealed with aluminum crimps. After filling and sealing, a 20 mL helium (He) headspace was introduced to each serum bottle, and the headspace was flushed with He twice for 3 min, with shaking in between. $O_2$ was then added back to reach *in situ* concentrations by injecting air using gas-tight syringes (Hamilton) with volumes calculated using the solubility equations of (2). One serum bottle per incubation depth contained optode sensor spots (Firesting, Pyroscience) for monitoring that air additions resulted in the desired *in situ* $O_2$ concentrations.

For nitrification and nitrite oxidation rate incubations, water was sampled directly from the Niskin bottle into 500 mL Tedlar bags (Restek) equipped with a septum injection port and three-way stopcocks for tracer addition and sampling, respectively (3). Bags were acid washed and purged with $N_2$ gas between incubations. Samples were drawn from each bag using a 60 mL syringe while applying constant pressure to the incubation bag to avoid air entering the bag. For depths where *in situ* $O_2$ concentrations were ≤5 µM, water was sampled into 120 mL glass serum bottles and a headspace was introduced as described above for DIC fixation rates.

#### *Nitrifier cultivation*

The ammonia-oxidizing bacterium *Nitrosomonas marina* C-25 and the nitrite-oxidizing bacterium *Nitrospina gracilis* Nb-3 were obtained from the culture collection of John B. Waterbury and Frederica Valois at the Woods Hole Oceanographic Institution (WHOI). The ammonia-oxidizing archaeon *Nitrosopumilus adriaticus* NF5 (=DSM 106890$^T$ = JCM 32270$^T$ = NCIMB 15114$^T$) is routinely grown in our labs and has been characterized in detail (4, 5).

*N. adriaticus* and *N. marina* were grown in HEPES-buffered artificial seawater medium containing 1 mM $NH_4Cl$ and 50 U $L^{-1}$ catalase (Cat. Nr. C9322; Sigma-Aldrich) as previously described (6). *N. gracilis* was grown in an artificial seawater medium supplemented with 1 mM $NaNO_2$ and 50 ng $L^{-1}$ cyanocobalamin (6). All strains were routinely grown in 60 mL polycarbonate bottles (Nalgene) containing 50 mL culture medium, and bottles were incubated at 25°C in the dark without agitation. To monitor culture growth, nitrite concentrations were measured using the Griess-Ilosvay colorimetric method (7).

***Library preparation and metagenome sequencing***

DNA libraries were prepared by the UC Davis Genome Center using the DNA Library Prep Kit with Fragmentation (Watchmaker Genomics, Boulder, CO) and Nextflex UDI sequencing adapters (Revvity) according to manufacturer's instructions. As the DNA extracts were partially denatured, a random priming protocol was used to generate homogeneously double-stranded DNA (8). The libraries were fragmented, amplified, analyzed via microcapillary electrophoresis on a LabChip GX Touch (Revvity), quantified by fluorometry on a Qubit instrument (LifeTechnologies), and pooled at equimolar concentrations. Pooled libraries were sequenced on two flowcells of the AVITI sequencer (Element Biosciences) run with paired-end 300 bp reads.

Prior to DNA extraction (described in the *Main Text*), DNA from three microbial genomes (*Deinococcus radiodurans* ATCC#13939, *Thermus thermophilus* ATCC#27634, *Blautia producta* ATCC#27340) was added as internal standards for downstream quantitative metagenomics (9). However, the development of quantitative metagenomics analysis methods was not within the scope of the current study. Consequently, internal standards were removed prior to metagenome analysis (methodological details in the *Main Text*).

***Metagenome-assembled-genomes and metagenomic read recruitment***

Metagenomic reads were trimmed, quality-filtered, and internal standards removed using established protocols (10, 11). Metagenomic sequences were assembled using MEGAHIT (v 1.2.9) (12) and metaSPAdes (v 3.14.1) (13). Assemblies were binned with metaWRAP (v 1.3.2) (14) using the binning tools MaxBin2 (v 2.2.6) (15), metaBAT2 (v 2.12.1) (16), and CONCOCT (v 1.0.0) (17). Bins completeness and contamination was estimated with CheckM (v 1.0.12) (18), and bins with a completeness >50% and contamination <10% were further refined using metaWRAP's "bin_refinement" module and dereplicated with dRep (v 3.5.0) (19). The resulting metagenome-assembled genomes (MAGs) were taxonomically classified using GTDB-Tk (v 2.3.2) (20).

We functionally annotated MAGs belonging to putative sulfur-oxidizing lineages. Open reading frames were predicted using Prodigal (v 2.6.3) (21) and predicted genes were annotated with eggNOG-mapper (v 2.1.12) (22–24). Putative chemoautotrophic sulfur oxidizers were identified by the presence of key genes in the Calvin-Benson-Bassam Cycle (CBB) and the reverse tricarboxylic acid (TCA) cycle, ribulose-1,5-bisphosphat-carboxylase/-oxygenase (RubisCO) and citrate lyase, respectively (Table S2). Metagenomic reads from each study site were mapped to MAGs of putative chemoautotrophic sulfur oxidizers using CoverM (v 0.7.0) (https://zenodo.org/records/10531254) with a minimum read percent identity of 95% and minimum read alignment of 75%.

### Supplementary Results and Discussion

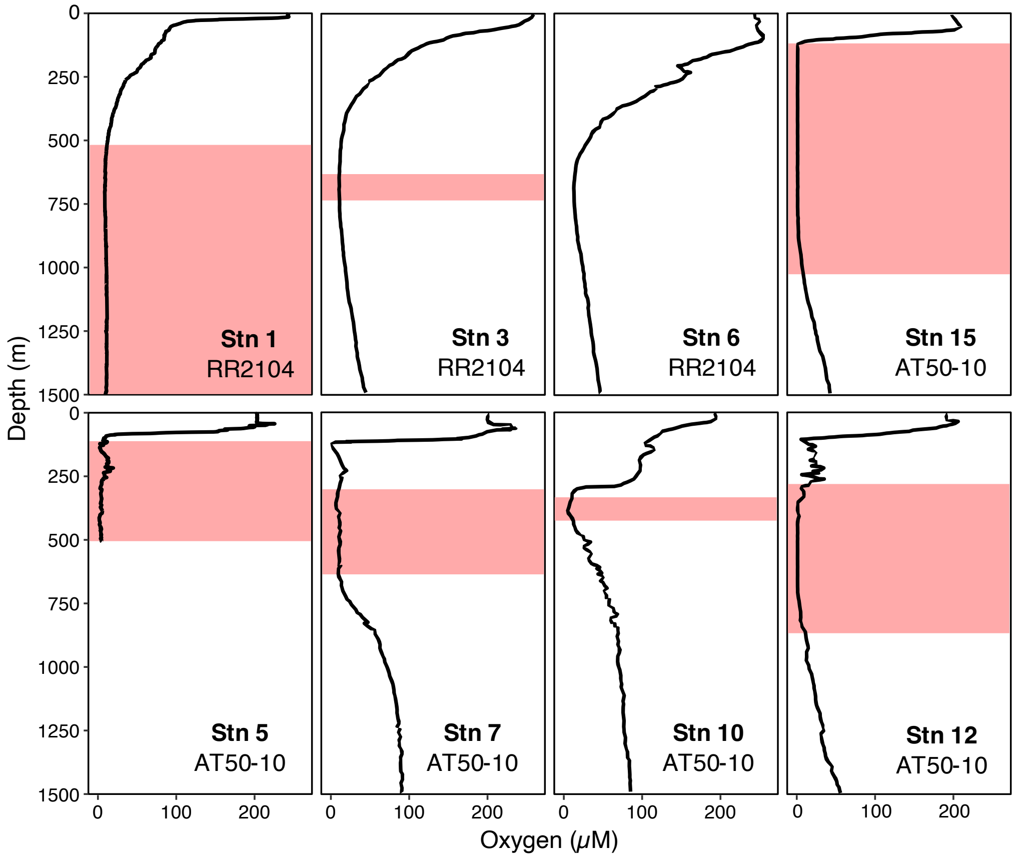


**Figure S1.** Oxygen concentration profiles of cruises RR2104 and AT50-10. The oxygen-deficient-zone (ODZ; $O_2 \leq 10$ $\mu$M) is shown by red shaded boxes. At Stn 6, $O_2$ concentrations never dropped below 12 $\mu$M.

***Relationships between nitrification rates and ammonia oxidizer abundances***

Nitrification rates in the open ocean are typically correlated with abundances of Water Column A (WCA) Clade AOA (*Ca.* Nitrosopelagicus) (Fig. S1, (3, 25)). Nitrification rates rapidly decrease with depth (*Main Text*, Fig. 1), while WCB Clade abundances increase with depth, explaining the negative correlation observed between both parameters (Fig. S2). In contrast to AOA of the WCA Clade and *Nitrosopumilus*, WCB Clade AOA only encode a high affinity version of an ammonium transporter (26), suggesting an exceptionally high substrate affinity in the deep ocean, where organic matter fluxes are low and ammonia concentrations undetectable. To date, no cultured representatives are available for WCB Clade AOA, yet all available metagenomic and single-cell genomic data indicate that they are indeed chemolithoautotrophic ammonia oxidizers (27, *and references therein*). Nevertheless, if or the extent to which they are also reliant on organic carbon compounds is currently not known. However, as turnover times for deep ocean AOA would be on the order of 1,500 days based on ammonia oxidation alone, there is a high chance that WCB AOA rely on other as-of-yet unknown metabolisms to support their abundance in the mesopelagic ocean (27, 28).

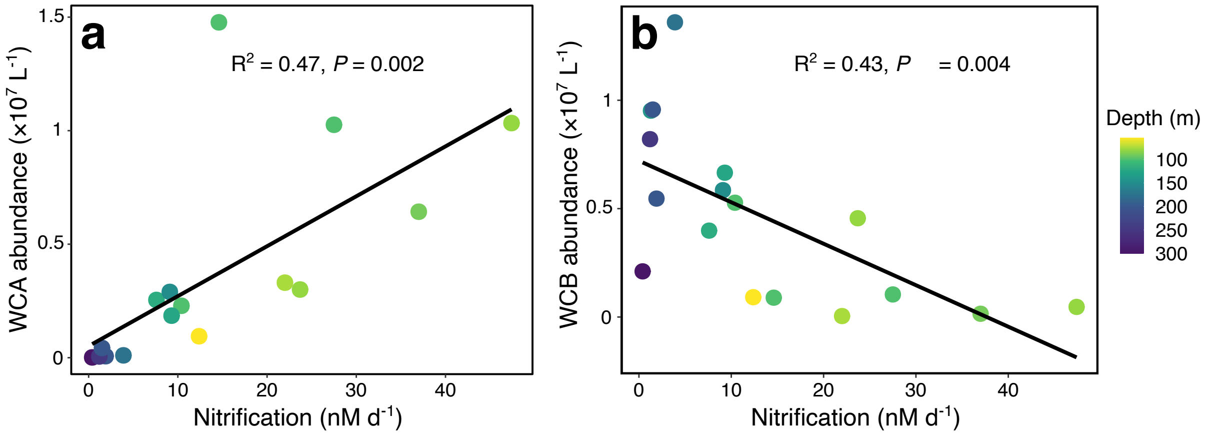


**Figure S2.** Relationships between nitrification rates and abundances of different clades of ammonia-oxidizing archaea during cruise AT50-10: (**a**) Water column A (WCA) Clade (**b**) Water column B (WCB) Clade. Sample depth is color-coded with darker colors reflecting deeper depths. Abundances were estimated from absolute *Nitrosopumilaceae* abundances derived from quantitative PCR assays and relative 16S rRNA gene abundances of each clade in metagenomes.

***Inhibition of marine nitrifier cultures by phenylacetylene***

As ammonia-oxidizing archaea (AOA) are the dominant ammonia oxidizers in most parts of the ocean, we determined the effective inhibitory concentration of phenylacetylene for *Nitrosopumilus adriaticus* NF5, which is one of the few available axenic cultures of marine AOA (5). Ammonia oxidation activity of *N. adriaticus* was completely inhibited by the addition of ≥ 5 $\mu$M phenylacetylene (Fig. S3a), which was lower than the inhibitory concentration determined for the terrestrial AOA *Nitrosocosmicus franklandicus* (≥20 $\mu$M) and within the same concentration range found to completely inhibit the terrestrial ammonia-oxidizing bacterium *Nitrosomonas europaea* (29). We selected a concentration of 10 $\mu$M phenylacetylene for all consecutive experiments and further confirmed that the DIC fixation activity of *N. adriaticus* and the marine ammonia-oxidizing bacterium *Nitrosomonas marina* was also inhibited after additions of 10 $\mu$M phenylacetylene (Fig. S3b). In contrast, the DIC fixation activity of the marine nitrite-oxidizing bacterium *Nitrospina gracilis* was not affected by phenylacetylene (Fig. S3b), in line with results from environmental nitrite oxidation rate experiments with and without phenylacetylene addition (*Main Text*, Fig. 3a), confirming that phenylacetylene only inhibits the first step in the nitrification process.

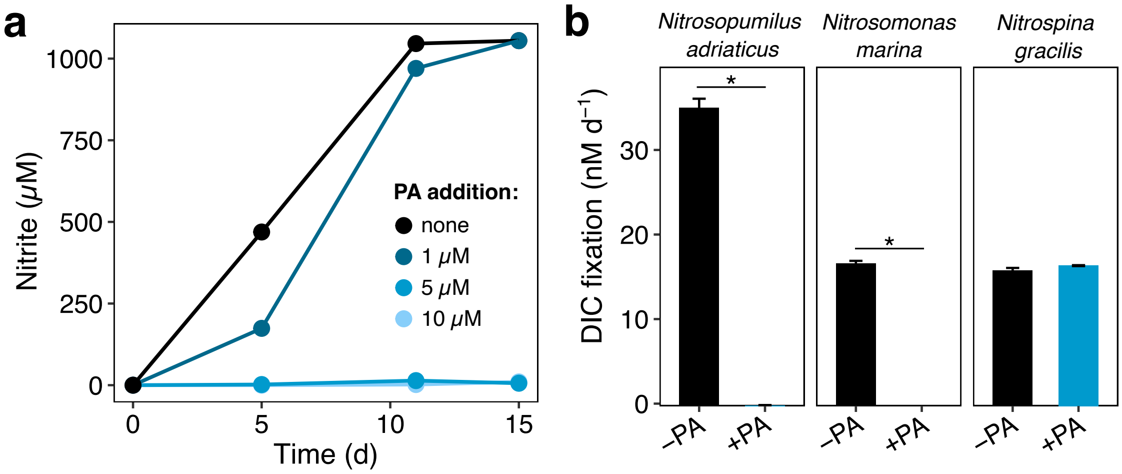


**Figure S3.** Effect of phenylacetylene on marine nitrifier cultures. **a)** Testing of different phenylacetylene (PA) concentrations on nitrite production (=ammonia-oxidizing activity) of *Nitrosopumilus adriaticus*. **b)** The effect of 10 $\mu$M phenylacetylene on DIC fixation rates of two marine ammonia oxidizers and the marine nitrite oxidizer *Nitrospina gracilis*. Error bars show the standard deviation of triplicate measurements. Significant differences between treatments (Student's *t*-test, $P < 0.05$) are indicated by an asterisk (*).

***Nitrite oxidation and nitrite oxidizer community composition***

Nitrite oxidation rates ranged from 3 to 45 nM $d^{-1}$ and typically followed nitrification rates (Fig S4 and *Main Text*, Fig. 3a), except for depths with low $O_2$ concentrations, where they were occasionally higher than nitrification rates as previously shown in the ETSP (3). In contrast, nitrification rates were sometimes higher than nitrite oxidation rates at shallow ( <100 m) depths (Fig. S4).

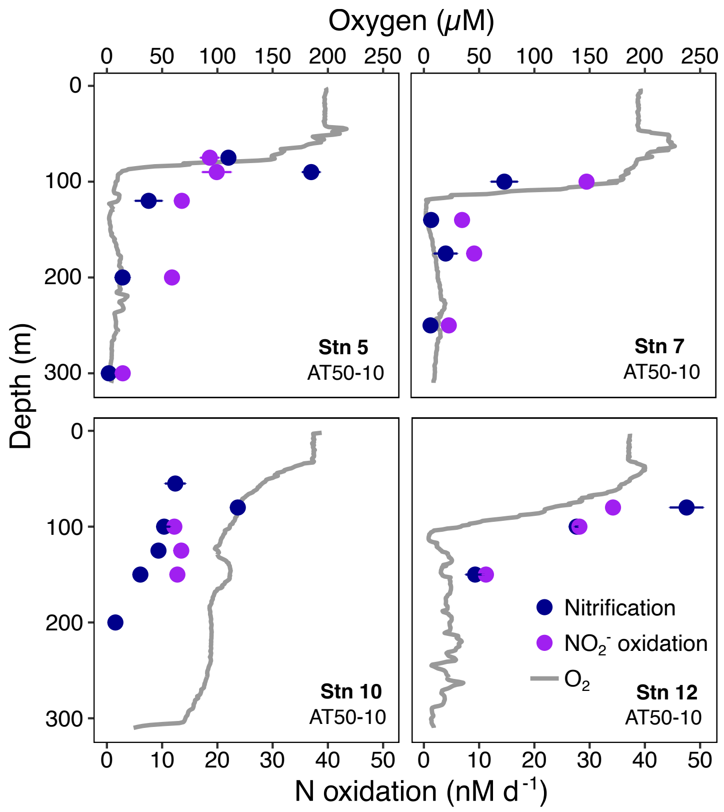


**Figure S4.** Nitrification and nitrite oxidation rates at four stations during cruise AT50-10. The mean and standard deviation of triplicate measurements are shown. Oxygen concentration profiles are depicted in gray.

Nitrite-oxidizing bacteria (NOB) of the *Nitrospinaceae* family were the main nitrite oxidizers at all stations and depths, making up 0.8-2.5% of the 16S rRNA gene sequences below the euphotic zone. *Nitrospinaceae* abundances were low in surface waters and sharply increased to ~$10^6$ cells $L^{-1}$ at the base of the euphotic zone (Fig. S5), coinciding with high abundances of AOA (*Main Text*, Fig. 2). Most sequences were affiliated with *Nitrospinaceae* Clade 1, while members of Clade 2 were lower in abundance but present at every station and almost every depth (Fig. S5). Additionally, 16S rRNA gene sequences affiliated to the genus *Nitrospira* were detected at low abundance (0.03-0.34%) at most depths at stations 5, 7 and 10.

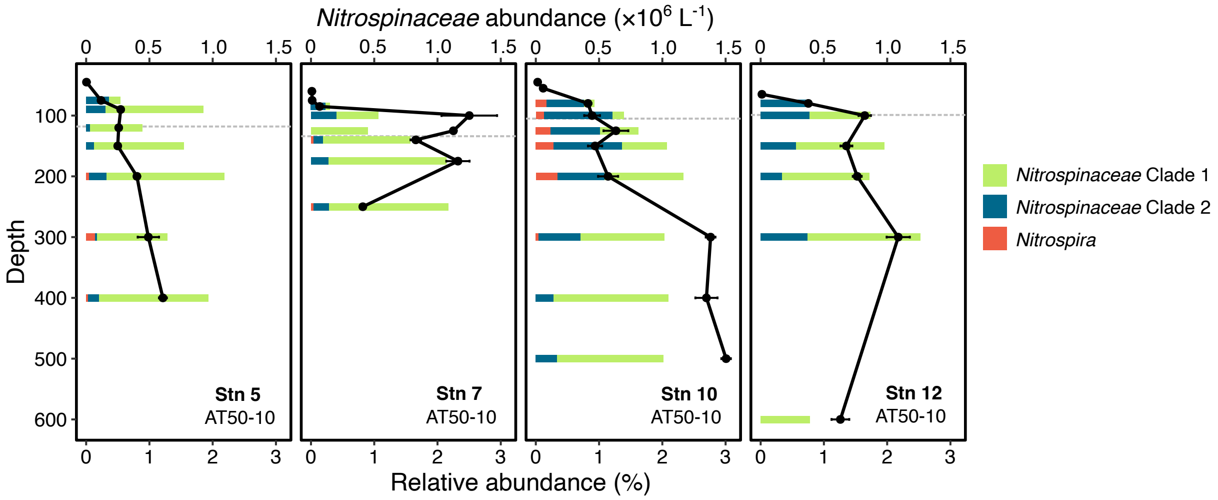


**Figure S5.** Nitrite oxidizer abundances and community composition at four stations of cruise AT50-10. Relative 16S rRNA gene abundances of different nitrite oxidizer clades are shown as a fraction of the total microbial community. Absolute *Nitrospinaceae* abundances were derived from quantitative PCR assays. Depth of the euphotic zone is indicated by gray dashed horizontal lines.

#### *Cell-specific DIC fixation rates of marine nitrifiers*

Estimated per-cell DIC fixation rates of AOA ranged from 0.02 to 0.36 fmol C cell$^{-1}$ d$^{-1}$ in the euphotic zone, which are within a similar range as rates reported for AOA cultures (0.04-0.82 fmol C cell$^{-1}$ d$^{-1}$, (6)), and slightly lower than AOA in the Gulf of Mexico (0.76 fmol C cell$^{-1}$ d$^{-1}$, (30)). In the mesopelagic, per-cell DIC fixation rate estimates were substantially lower, ranging from 0.002 to 0.08 fmol C cell$^{-1}$ d$^{-1}$, as previously reported for North Atlantic deep waters (0.002-0.1 fmol C cell$^{-1}$ d$^{-1}$, (31)).

Using DIC fixation yields established for cultured *Nitrospina* (6), estimated cell-specific DIC fixation rates of *Nitrospinaceae* ranged from 0.23 to 6 fmol cell$^{-1}$ d$^{-1}$, which is higher than reported in the North Atlantic (0.002-0.734 fmol cell$^{-1}$ d$^{-1}$, (32)) and for cultured *Nitrospina* (0.17-1.54 fmol cell$^{-1}$ d$^{-1}$, (6)). This discrepancy could be explained by lower DIC fixation yields of nitrite oxidizers in the environment compared to yields obtained from cultures. Alternatively, cell-specific DIC fixation rates of *Nitrospinaceae* could be higher in regions with low $O_2$ concentrations, in contrast to those inhabiting fully oxygenated waters of the North Atlantic or grown under controlled culture conditions.

***Putative chemolithoautotrophic sulfur oxidizers***

Chemolithoautotrophic sulfur oxidizers in the dark ocean are phylogenetically diverse, including representatives of the SAR324 clade, *Marinisomataceae* (SAR406 lineage), *Thioglobaceae* (SUP05 lineage) and UBA868 family (33–36). *Marinisomataceae*, *Thioglobaceae* and members of the SAR324 clade were abundant at all stations, together making up to 28% of the 16S rRNA gene sequences in the mesopelagic (Fig. S6a), comparable to the abundance of AOA (*Main Text,* Fig. 2). However, each of these lineages also contain heterotrophic representatives (33, 35, 37, 38) making a direct connection between the abundance of these groups to DIC fixation rates challenging. Indeed, many of the metagenome-assembled-genomes (MAGs) of these phylogenetic groups recovered from our study sites were lacking genes involved in autotrophic DIC fixation pathways (Table S4). Overall, 17% of the *Marinisomataceae* and 67% of the *Thioglobaceae* MAGs encoded genes for ribulose-1,5-bisphosphat-carboxylase/-oxygenase (RubisCO), while 29% of the MAGs of the SAR324 clade encoded for citrate lyase, key genes in the Calvin-Benson-Bassam Cycle (CBB) and the reverse tricarboxylic acid (TCA) cycle, respectively (Table S4). Additionally, we recovered ten MAGs of the newly discovered gammaproteobacterial sulfur oxidizers of the UBA868 clade (36) (which could not be identified via 16S rRNA gene classifications), of which five encoded RubisCO genes.

We mapped metagenomic reads to MAGs encoding DIC fixation genes to estimate the relative abundance of putative autotrophic sulfur oxidizers at our study sites (Fig. S6b). Compared to the 16S rRNA gene-based relative abundances of phylogenetic groups containing sulfur oxidizers, the relative abundances of MAGs encoding DIC fixation genes were ~10 times lower, suggesting a dominance of heterotrophic representatives of these phylogenetic groups (Fig. S6). While these results can be biased by the presumed absence of genes due to the incompleteness of MAGs (>50% completeness), such large discrepancies are unlikely to result from MAG incompleteness alone. Chemolithoautotrophic *Thioglobaceae* and *Marinisomataceae* had the highest relative abundance below the euphotic zone (~1-2.5% and ~0.2%, respectively), while UBA868 MAGs exhibited a higher relative abundance within the euphotic zone (≤1%), except for the anoxic core at Stn 12 (600 m depth), where UBA868 and *Marinisomataceae* abundances were highest (Fig. S6b). No reads could be mapped to chemolithoautotrophic SAR324 MAGs, suggesting that members of the SAR324 clade at the investigated stations and depths might have a heterotrophic lifestyle.

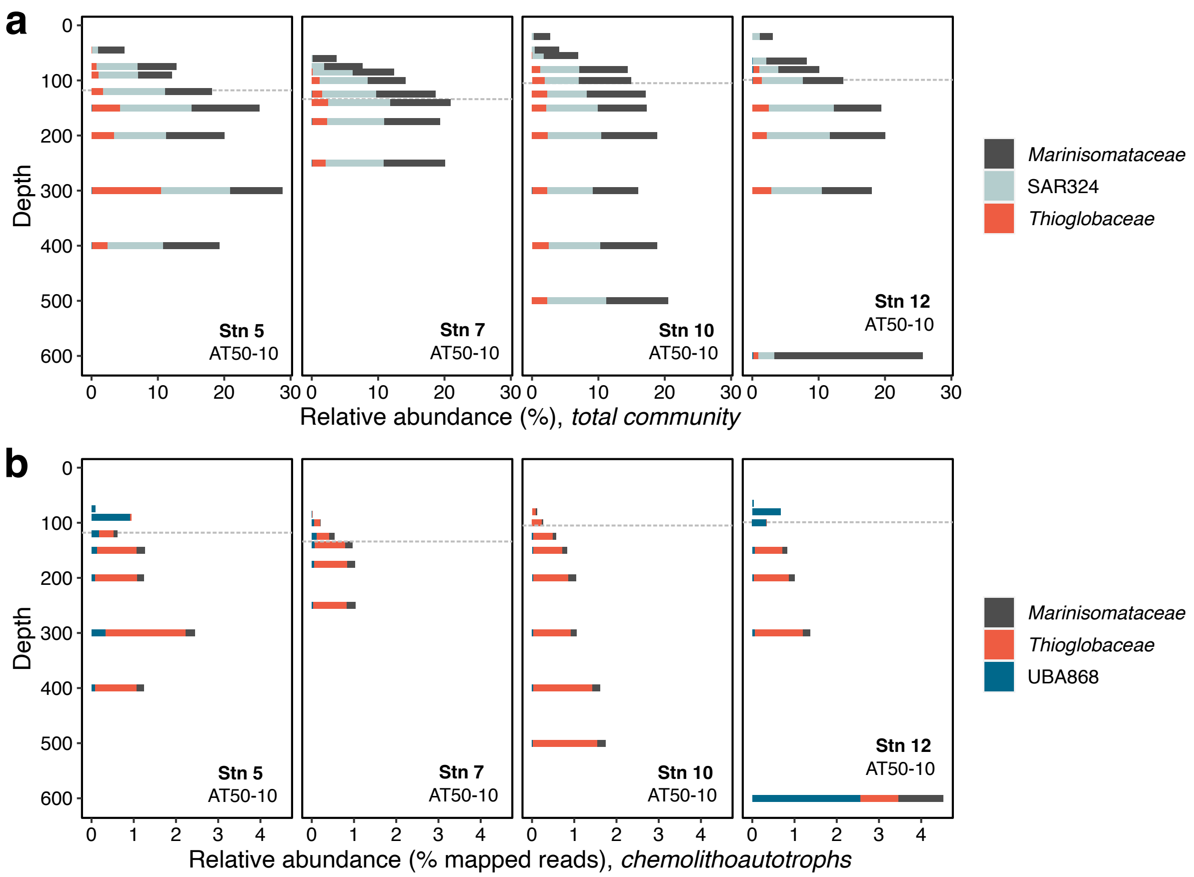


**Figure S6.** Relative abundances of phylogenetic groups containing chemolithoautotrophic sulfur oxidizers during cruise AT50-10 based on (**a**) 16S rRNA gene sequences affiliated with each phylogenetic group (*total community*), and (**b**) metagenomic reads mapped to MAGs containing DIC fixation genes (*chemolithoautotrophs*). Depth of the euphotic zone is indicated by gray dashed horizontal lines.

***Presumed stimulation of dark DIC fixation by plastic leachates***

At the start of cruise AT50-10, we performed $^{14}C$-bicarbonate incorporation experiments from oxic depths in 50 mL conical centrifuge tubes (Fisher Scientific). These tubes are made from polypropylene, are commonly used for diverse laborarory applications, and are practical for disposal when contaminated with radioactivity. However, we noticed high and very variable dark DIC fixation rates when seawater was incubated in plastic tubes (Fig. S7). Dark DIC fixation rates were on average 2.2-5.4 times higher in polypropylene compared to glass tubes, and acid-washing of these tubes had no effect (Fig. S7). Consequently, we switched to 40 mL glass vials with teflon coated silicon septa (TOC-certified, Fisher Scientific) after evaluating the results of the first stations. We hypothesize that plastic leachates stimulated dark DIC fixation by heterotrophic microbes. The types and specifics of different plastic additives (such as plasticizers) are typically not provided by manufacturers, making it difficult to pinpoint which substances stimulated dark DIC fixation in our study. However, we suggest caution is warranted in using plastic tubes for DIC fixation rate measurements in marine environments. Differences among plastic tube brands have previously been observed for leucine incorporation rates (39), however, these were largely attributed to differences in protein retention, which is not a concern when biomass is retained on membrane filters.

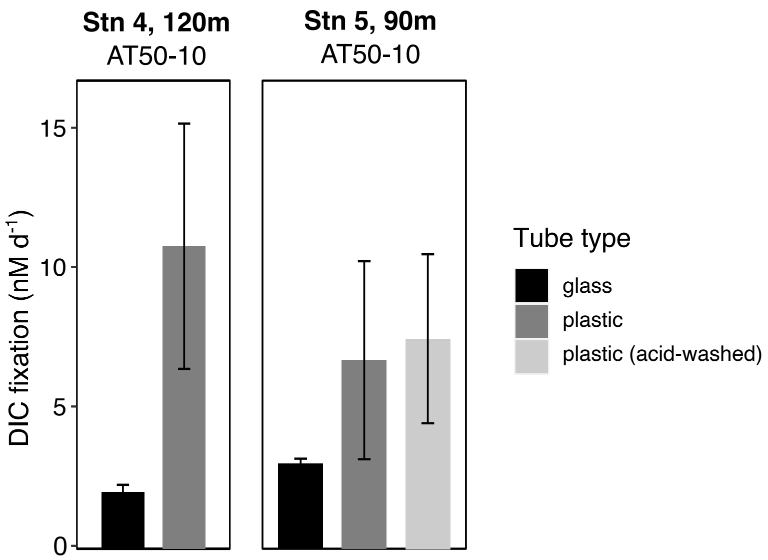


**Figure S7.** Comparison between dark DIC fixation rates measured in plastic *vs*. glass tubes. Shown are the mean and standard deviation of triplicate measurements.

***Comparison of dark DIC fixation rates obtained from $^{13}C$ vs. $^{14}C$ incorporation***

Due to restrictions in using radioactive materials on cruise RR2104, we used [$^{13}C$]-labeled bicarbonate to assess dark DIC fixation rates. In contrast, [$^{14}C$]-labeled bicarbonate was used to measure dark DIC fixation rates on cruise AT50-10 (methodological details in the *Main Text*). To verify comparability between rates obtained from both cruises, we used the ammonia-oxidizing archaeon *Nitrosopumilus adriaticus* to compare DIC fixation rates measured by [$^{14}C$] vs [$^{13}C$]-bicarbonate incorporation. We found a good agreement between the two methods, with slightly higher but statistically not significantly different (Student's *t*-test, *P* value=0.097) DIC fixation rates from [$^{13}C$]-bicarbonate (mean±sd=2.1±0.1 $\mu$M d$^{-1}$) compared to [$^{14}C$]-bicarbonate incorporation (mean±sd=1.9±0.1 $\mu$M d$^{-1}$) (Fig S8a).

Due to the small volumes (40 mL) used for [$^{14}C$]-bicarbonate assimilation assays and low microbial activities in the dark ocean, we used long (48-72 h) incubation times to ensure sufficient radiocarbon incorporation. We analyzed the linearity of dark DIC fixation rates at a few select stations and depths and found that rates were always linear (Fig. S8b), suggesting no stimulation of dark DIC fixation over the length of the incubation in our study.

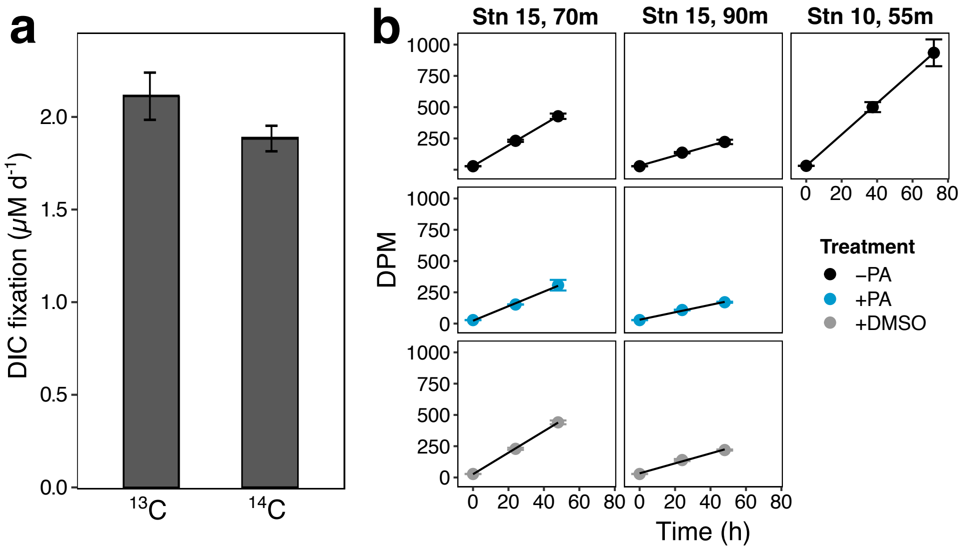


**Figure S8.** Method validation of dark DIC fixation rates. **a**) Comparison of [$^{13}C$] *vs*. [$^{14}C$]-bicarbonate incorporation by the ammonia-oxidizing archaeon *Nitrosopumilus adriaticus*. **b**) Linearity of dark DIC fixation rates over 2-3 days from [$^{14}C$]-bicarbonate incorporation experiments performed during cruise AT50-10. Shown are the mean and standard deviation of triplicate measurements.

**Table S1**. Contributions of different metabolisms to depth-integrated dark DIC fixation rates in the eastern tropical and subtropical Pacific Ocean. Total and ammonia-fueled dark DIC fixation rates are measured directly, while contributions of other metabolisms were estimated (details in the *Main Text*). Estimates of nitrite-fueled chemoautotrophy are based on measured nitrite oxidation rates and DIC fixation yields of nitrite oxidizer cultures (6). Heterotrophic DIC fixation rates were estimated as 10% of measured heterotrophic production rates. Sulfur-fueled chemoautotrophy was estimated based on differences in the N:S stoichiometry of particulate organic matter (40), and differences between the DIC fixation yields of nitrifiers (*Main Text,* Fig. 5a, and (6)) and sulfide oxidizers (41–43).

| **Depth-integrated rates ($\mu$mol m$^{-2}$ d$^{-1}$)** | **Station 5** | **Station 7** | **Station 10** | **Station 12** |
|---|---|---|---|---|
| Total dark DIC fixation | 215 | 256 | 328 | 849 |
| Ammonia-fueled chemoautotrophy | 48 | 5 | 55 | 165 |
| Nitrite-fueled chemoautotrophy | 82 | 76 | 23 | 57 |
| Sulfur-fueled chemoautotrophy | 19* / 47# | 2* / 43# | 22* / 13# | 68* / 33# |
| Heterotrophic DIC fixation | 95 | 99 | 107 | 203 |

*Calculated based on estimates of ammonia-fueled chemoautotrophy.

#Calculated based on estimates of nitrite-fueled chemoautotrophy.

**Table S2.** List of metagenome-assembled-genomes (MAGs) from taxonomic groups containing putative chemolithoautotrophic sulfur oxidizers.

| MAG | RubisCO | Citrate lyase | Completeness (%) | Contamination (%) | Sample name | SRA Accessions |
|---|---|---|---|---|---|---|
| **SAR324** | | | | | | |
| CLZ33_megahit_bin.12 | X | | 51.18 | 1.68 | CLZ33 | SRR31341480, SRR31341479 |
| CLZ125_metaspades_bin.10 | | | 90.14 | 0 | CLZ125 | SRR31341472, SRR31341471 |
| CLZ219_megahit_bin.12 | | | 92.3 | 2.323 | CLZ219 | SRR31156210, SRR31156209 |
| CLZ219_metaspades_bin.12 | | | 51.37 | 3.588 | CLZ219 | SRR31156210, SRR31156209 |
| CLZ221_metaspades_bin.2 | | | 77.66 | 1.148 | CLZ221 | SRR31156213, SRR31156211 |
| CLZ223_megahit_bin.2 | | | 50.6 | 7.3 | CLZ223 | SRR31156214, SRR31156215 |
| CLZ233_megahit_bin.13 | X | | 60.53 | 9.248 | CLZ233 | SRR31341478, SRR31341477 |
| **UBA868** | | | | | | |
| CLZ65_megahit_bin.11 | X | | 54.12 | 6.504 | CLZ65 | SRR31156262, SRR31156263 |
| CLZ75_megahit_bin.1 | X | | 92.92 | 3.14 | CLZ75 | SRR31156217, RR31156234 |
| CLZ129_metaspades_bin.1 | | | 70.18 | 1.219 | CLZ129 | SRR31341473, SRR31341476 |
| CLZ181_metaspades_bin.4 | | | 62.01 | 3.618 | CLZ181 | SRR31156235, SRR31156236 |
| CLZ211_metaspades_bin.15 | X | | 82.77 | 3.252 | CLZ211 | SRR31156226, SRR31156227 |
| CLZ211_metaspades_bin.19 | X | | 95.02 | 1.829 | CLZ211 | SRR31156226, SRR31156227 |
| CLZ215_metaspades_bin.1 | | | 96.64 | 2.439 | CLZ215 | SRR31156205, SRR31156206 |
| CLZ219_megahit_bin.9 | | | 76.52 | 1.629 | CLZ219 | SRR31156210, SRR31156209 |
| CLZ219_metaspades_bin.1 | X | | 56.57 | 1.036 | CLZ219 | SRR31156210, SRR31156209 |
| CLZ233_megahit_bin.2 | | | 93.59 | 3.048 | CLZ233 | SRR31341478, SRR31341477 |
| ***Thioglobaceae* (SUP05)** | | | | | | |
| CLZ57_metaspades_bin.5 | | | 51.74 | 5.573 | CLZ57 | SRR31156261, SRR31156264 |
| CLZ65_megahit_bin.8 | X | | 55.66 | 0.993 | CLZ65 | SRR31156262, SRR31156263 |
| CLZ211_metaspades_bin.11 | X | | 94.77 | 2.851 | CLZ211 | SRR31156226, SRR31156227 |
| CLZ219_megahit_bin.1 | | | 92.3 | 2.323 | CLZ219 | SRR31156210, SRR31156209 |
| CLZ233_metaspades_bin.2 | X | | 68.16 | 0 | CLZ233 | SRR31341478, SRR31341477 |
| CLZ237_metaspades_bin.8 | X | | 58.55 | 1.379 | CLZ237 | SRR31341475, SRR31341474 |
| ***Marinisomataceae* (SAR406)** | | | | | | |
| CLZ65_megahit_bin.9 | | | 75.46 | 9.89 | CLZ65 | SRR31156262, SRR31156263 |
| CLZ77_metaspades_bin.1 | | | 59.89 | 1.098 | CLZ77 | SRR31156255, SRR31156248 |
| CLZ83_megahit_bin.7 | | | 67.45 | 1.098 | CLZ83 | SRR31341469, SRR31341470 |

| | | | | | | |
|---|---|---|---|---|---|---|
| CLZ89_metaspades_bin.5 | | | 51.72 | 1.724 | CLZ89 | SRR31341466, SRR31341467 |
| CLZ89_metaspades_bin.6 | | | 52.85 | 8.306 | CLZ89 | SRR31341466, SRR31341467 |
| CLZ127_metaspades_bin.4 | | | 52.12 | 2.197 | CLZ127 | SRR31341465, SRR31341468 |
| CLZ155_megahit_bin.8 | | | 52.98 | 0 | CLZ155 | SRR31341463, SRR31341464 |
| CLZ159_metaspades_bin.1 | | | 69.78 | 0.099 | CLZ159 | SRR31156223, SRR31156221 |
| CLZ171_megahit_bin.4 | | X | 50.92 | 9.89 | CLZ171 | SRR31156228, SRR31156229 |
| CLZ181_megahit_bin.2 | | | 54.69 | 2.791 | CLZ181 | SRR31156235, SRR31156236 |
| CLZ211_metaspades_bin.2 | | | 69.31 | 2.991 | CLZ211 | SRR31156226, SRR31156227 |
| CLZ211_metaspades_bin.6 | | | 91.14 | 0 | CLZ211 | SRR31156226, SRR31156227 |
| CLZ211_metaspades_bin.7 | | | 86.81 | 2.197 | CLZ211 | SRR31156226, SRR31156227 |
| CLZ211_metaspades_bin.9 | | X | 83.45 | 0 | CLZ211 | SRR31156226, SRR31156227 |
| CLZ219_metaspades_bin.5 | | | 89.81 | 1.648 | CLZ219 | SRR31156210, SRR31156209 |
| CLZ233_metaspades_bin.5 | | | 87.61 | 0 | CLZ233 | SRR31341478, SRR31341477 |
| CLZ233_metaspades_bin.10 | | X | 58.23 | 1.098 | CLZ233 | SRR31341478, SRR31341477 |

### Supplementary References


1. L. De Brabandere, B. Thamdrup, N. P. Revsbech, R. Foadi, A critical assessment of the occurrence and extend of oxygen contamination during anaerobic incubations utilizing commercially available vials. *J. Microbiol. Methods* **88**, 147–154 (2012).

2. H. E. Garcia, L. I. Gordon, Oxygen solubility in seawater: Better fitting equations. *Limnol. Oceanogr.* **37**, 1307–1312 (1992).

3. A. E. Santoro, *et al.*, Nitrification and nitrous oxide production in the offshore waters of the Eastern Tropical South Pacific. *Glob. Biogeochem. Cycles* **35**, 0e2020GB006716-3 (2021).

4. B. Bayer, *et al.*, Physiological and genomic characterization of two novel marine thaumarchaeal strains indicates niche differentiation. *ISME J.* **10**, 1051–1063 (2016).

5. B. Bayer, *et al.*, Nitrosopumilus adriaticus sp. nov. and Nitrosopumilus piranensis sp. nov., two ammonia-oxidizing archaea from the Adriatic Sea and members of the class Nitrososphaeria. *Int. J. Syst. Evol. Microbiol.* **7**, 1892–1902 (2019).

6. B. Bayer, K. McBeain, C. A. Carlson, A. E. Santoro, Carbon content, carbon fixation yield and dissolved organic carbon release from diverse marine nitrifiers. *Limnol. Oceanogr.* **68**, 84–96 (2023).

7. J. D. H. Strickland, T. R. Parsons, A Partical Handbook of Seawater Analysis. *Fish Res Bd Can Bull No 167* (1972).

8. E. A. Saunderson, *et al.*, A novel use of random priming-based single-strand library preparation for whole genome sequencing of formalin-fixed paraffin-embedded tissue samples. *NAR Genomics Bioinforma.* **2**, lqz017 (2020).

9. S. M. Gifford, *et al.*, Microbial Niche Diversification in the Galápagos Archipelago and Its Response to El Niño. *Front. Microbiol.* **11**, 575194 (2020).

10. B. Langmead, S. L. Salzberg, Fast gapped-read alignment with Bowtie 2. *Nat. Methods* **9**, 357–359 (2012).

11. A. M. Bolger, M. Lohse, B. Usadel, Trimmomatic: a flexible trimmer for Illumina sequence data. *Bioinformatics* **30**, 2114–2120 (2014).

12. D. Li, C.-M. Liu, R. Luo, K. Sadakane, T.-W. Lam, MEGAHIT: an ultra-fast single-node solution for large and complex metagenomics assembly via succinct *de Bruijn* graph. *Bioinformatics* **31**, 1674–1676 (2015).

13. A. Prjibelski, D. Antipov, D. Meleshko, A. Lapidus, A. Korobeynikov, Using SPAdes De Novo Assembler. *Curr. Protoc. Bioinforma.* **70**, e102 (2020).

14. G. V. Uritskiy, J. DiRuggiero, J. Taylor, MetaWRAP—a flexible pipeline for genome-resolved metagenomic data analysis. *Microbiome* **6**, 158 (2018).

15. Y.-W. Wu, B. A. Simmons, S. W. Singer, MaxBin 2.0: an automated binning algorithm to recover genomes from multiple metagenomic datasets. *Bioinformatics* **32**, 605–607 (2016).

16. D. D. Kang, J. Froula, R. Egan, Z. Wang, MetaBAT, an efficient tool for accurately reconstructing single genomes from complex microbial communities. *PeerJ* **3**, e1165 (2015).

17. J. Alneberg, *et al.*, Binning metagenomic contigs by coverage and composition. *Nat. Methods* **11**, 1144–1146 (2014).

18. D. H. Parks, M. Imelfort, C. T. Skennerton, P. Hugenholtz, G. W. Tyson, CheckM: Assessing the quality of microbial genomes recovered from isolates, single cells, and metagenomes. *Genome Res.* **25**, 1043–1055 (2015).

19. M. R. Olm, C. T. Brown, B. Brooks, J. F. Banfield, dRep: a tool for fast and accurate genomic comparisons that enables improved genome recovery from metagenomes through de-replication. *ISME J.* **11**, 2864–2868 (2017).

20. P.-A. Chaumeil, A. J. Mussig, P. Hugenholtz, D. H. Parks, GTDB-Tk: a toolkit to classify genomes with the Genome Taxonomy Database. *Bioinformatics* **36**, 1925–1927 (2019).

21. D. Hyatt, *et al.*, Prodigal: prokaryotic gene recognition and translation initiation site identification. *BMC Bioinformatics* **11**, 119 (2010).

22. C. P. Cantalapiedra, A. Hernández-Plaza, I. Letunic, P. Bork, J. Huerta-Cepas, eggNOG-mapper v2: Functional Annotation, Orthology Assignments, and Domain Prediction at the Metagenomic Scale. *Mol. Biol. Evol.* **38**, 5825–5829 (2021).

23. J. Huerta-Cepas, *et al.*, eggNOG 5.0: a hierarchical, functionally and phylogenetically annotated orthology resource based on 5090 organisms and 2502 viruses. *Nucleic Acids Res.* **47**, D309–D314 (2019).

24. M. Steinegger, J. Söding, MMseqs2 enables sensitive protein sequence searching for the analysis of massive data sets. *Nat. Biotechnol.* **35**, 1026–1028 (2017).

25. A. E. Santoro, B. Bayer, F. J. Elling, A. Pearson, "Candidatus Nitrosopelagicus" in *Bergey's Manual of Systematics of Archaea and Bacteria*, (2021), pp. 1–13.

26. W. Qin, *et al.*, Alternative strategies of nutrient acquisition and energy conservation map to the biogeography of marine ammonia-oxidizing archaea. *ISME J.* **14**, 2595–2609 (2020).

27. A. E. Santoro, R. A. Richter, C. L. Dupont, Planktonic Marine Archaea. *Annu. Rev. Mar. Sci.* **11**, 131–158 (2019).

28. S. E. Newell, A. R. Babbin, A. Jayakumar, B. B. Ward, Ammonia oxidation rates and nitrification in the Arabian Sea. *Glob. Biogeochem. Cycles* **25**, n/a-n/a (2011).

29. C. L. Wright, A. Schatteman, A. T. Crombie, J. C. Murrell, L. E. Lehtovirta-Morley, Inhibition of Ammonia Monooxygenase from Ammonia-Oxidizing Archaea by Linear and Aromatic Alkynes. *Appl. Environ. Microbiol.* **86**, e02388-19 (2020).

30. K. Kitzinger, *et al.*, Single cell analyses reveal contrasting life strategies of the two main nitrifiers in the ocean. *Nat. Commun.* **11** (2020).

31. M. M. Varela, H. M. van Aken, E. Sintes, T. Reinthaler, G. J. Herndl, Contribution of Crenarchaeota and Bacteria to autotrophy in the North Atlantic interior. *Environ. Microbiol.* **13**, 1524–1533 (2011).

32. M. G. Pachiadaki, *et al.*, Major role of nitrite-oxidizing bacteria in dark ocean carbon fixation. *Science* **358**, 1046–1051 (2017).

33. A. K. Hawley, *et al.*, Diverse Marinimicrobia bacteria may mediate coupled biogeochemical cycles along eco-thermodynamic gradients. *Nat. Commun.* **8** (2017).

34. L. Malfertheiner, C. Martínez-Pérez, Z. Zhao, G. J. Herndl, F. Baltar, Phylogeny and Metabolic Potential of the Candidate Phylum SAR324. *Biology* **11**, 599 (2022).

35. R. M. Morris, R. L. Spietz, The Physiology and Biogeochemistry of SUP05. *Annu. Rev. Mar. Sci.* **14**, 261–275 (2022).

36. F. Baltar, *et al.*, A ubiquitous gammaproteobacterial clade dominates expression of sulfur oxidation genes across the mesopelagic ocean. *Nat. Microbiol.* **8**, 1137–1148 (2023).

37. R. L. Spietz, *et al.*, Heterotrophic carbon metabolism and energy acquisition in Candidatus Thioglobus singularis strain PS1, a member of the SUP05 clade of marine Gammaproteobacteria. *Environ. Microbiol.* **21**, 2391–2401 (2019).

38. D. Boeuf, *et al.*, Metapangenomics reveals depth-dependent shifts in metabolic potential for the ubiquitous marine bacterial SAR324 lineage. *Microbiome* **9** (2021).

39. M. L. Pace, P. Del Giorgio, D. Fischer, R. Condon, H. Malcom, Estimates of bacterial production using the leucine incorporation method are influenced by differences in protein retention of microcentrifuge tubes. *Limnol. Oceanogr. Methods* **2**, 55–61 (2004).

40. P. A. Matrai, R. W. Eppley, Particulate organic sulfur in the waters of the Southern California Bight. *Glob. Biogeochem. Cycles* **3**, 89–103 (1989).

41. D. C. Nelson, B. B. Jørgensen, N. P. Revsbech, Growth Pattern and Yield of a Chemoautotrophic *Beggiatoa* sp. in Oxygen-Sulfide Microgradients. *Appl. Environ. Microbiol.* **52**, 225–233 (1986).

42. J. M. Klatt, L. Polerecky, Assessment of the stoichiometry and efficiency of $CO_2$ fixation coupled to reduced sulfur oxidation. *Front. Microbiol.* **6** (2015).

43. D. Vasquez-Cardenas, F. J. R. Meysman, H. T. S. Boschker, A Cross-System Comparison of Dark Carbon Fixation in Coastal Sediments. *Glob. Biogeochem. Cycles* **34**, e2019GB006298 (2020).
